# Situating affective, autonomic, and wakefulness arousal in a connectome-based neural reference space

**DOI:** 10.64898/2026.06.08.730730

**Authors:** Kannon Bhattacharyya, Judah Huberman-Shlaes, Yulin Tong, Yuxuan Zhu, Jin Ke, Jadyn S. Park, Anna Corriveau, Monica D. Rosenberg, Yuan Chang Leong

**Author notes:** Correspondence should be addressed to Yuan Chang Leong. These authors contributed equally (Kannon Bhattacharyya, Judah Huberman-Shlaes).

## Abstract

Arousal is often invoked to explain state-dependent variability in attention, emotion, memory, and decision-making. However, the term is used inconsistently, referring variously to affective experience, autonomic activation, and states of wakefulness. The extent to which different operationalizations of arousal converge remains unclear, limiting efforts to integrate findings across studies. Here, we situate different varieties of arousal within a connectome-based reference space, assessing their convergence through predictive generalization across measures and the similarity of their predictive connectivity patterns. We applied dynamic connectome-based predictive models to five fMRI datasets spanning naturalistic movie watching, story listening, wakeful rest, and sleep. We derived measures of affective, autonomic, and wakefulness arousal from subjective ratings, pupil dilation, and EEG, respectively. Models trained to predict arousal in one dataset generalized across datasets, indicating that dynamic connectivity captures shared arousal-related dynamics across measures and task contexts. Predictive connections partially overlapped across models, including a subset shared across all four models. The models, as well as a model restricted to the shared connections, predicted manually scored sleep stages, suggesting that they capture arousal dynamics that extend to the graded fluctuations in arousal during sleep. Decoded arousal dynamics during movie-viewing predicted how well participants later recalled movie events, reproducing classic arousal-dependent memory enhancement effects. Together, these results suggest that different measures of arousal partially overlap within a connectome-based reference space, sharing connectivity-predictable variance that is behaviorally relevant. More broadly, this approach offers an empirical framework for assessing how different operationalizations of arousal relate to one another, and can be extended to other measures, populations, and psychological states.

## Introduction

The quickened pulse before an exam, the restless wakefulness in the middle of the night, and the sharpened focus in a high-stakes moment are thought to reflect fluctuations in arousal. Broadly defined as a state of heightened activation, arousal has been proposed as a core explanatory factor across models of attention [1–4], emotion [5,6], learning [7,8], memory [9–12], and decision-making [13–16]. Yet, what counts as “arousal” varies considerably across studies and theories [17]. In some contexts, the term refers to a subjective experience, such as feeling excited, anxious, or emotionally charged [18,19]. In others, it refers to autonomic nervous system activity, including changes in heart rate, pupil dilation, or skin conductance [20–23]. Arousal has also been used to describe states of wakefulness, spanning anesthesia, sleep, drowsiness, and full alertness [24–26]. During emotional episodes, subjective experience, physiology, and behavior are only modestly correlated [27–29], and even different physiological measures do not always agree [23,30,31]. This heterogeneity, both within and between varieties, has led critics to question the construct’s scientific utility, arguing that collapsing distinct processes under a single label obscures mechanistic specificity and hinders theoretical progress [32–35].

In light of these critiques, a reasonable response would be to abandon the umbrella term “arousal” altogether in favor of more specific constructs (e.g., affective intensity, autonomic activation, vigilance). However, this risks obscuring potential commonalities across varieties of arousal and discarding the cumulative insights gained from decades of arousal research. Indeed, the persistent use of the term suggests an intuitive recognition that these phenomena may be meaningfully related. Rather than replacing arousal with a set of narrower terms, we take a complementary approach and ask whether common structure can be empirically identified across different types of arousal. We draw on the concept of a *neural reference space* [36], conceptualized as the set of neural responses probabilistically associated with a class of mental events, with different instances potentially engaging different, partially overlapping patterns. A neural reference space of arousal would thus define a shared space in which different operationalizations of arousal can be situated and their convergence assessed empirically.

Recent efforts have begun to construct a neural reference space of arousal using comparative reviews [17] and meta-analytic approaches [37,38]. These studies have identified overlapping involvement of brainstem nuclei, including the locus coeruleus and dorsal raphe nucleus, the thalamus, and regions in the salience network, such as the anterior insula and dorsal anterior cingulate cortex. However, regional overlap does not guarantee that the same neural states underlie different varieties of arousal. A given region can support multiple functions depending on which neural populations are engaged and their pattern of connectivity with other regions. Indeed, a recent study [39] derived a distributed pattern of brain activity that predicts affective arousal across diverse tasks and stimuli. Although this pattern showed spatial overlap with many of the regions highlighted in prior meta-analytic maps of arousal, it did not generalize to predict autonomic activation or states of wakefulness.

Furthermore, studies that have empirically characterized neural reference spaces of arousal have primarily relied on activity patterns rather than interactions among brain regions [39,40]. Yet a growing literature indicates that arousal is closely tied to changes in functional connectivity [41–49]. For example, autonomic activation has been associated with increased connectivity within the salience network [45], as well as stronger coupling between salience network regions and the thalamus, brainstem, and dorsolateral prefrontal cortex [44]. Fluctuations in affective arousal have likewise been shown to covary with dynamic connectivity among large-scale functional networks, including salience, default mode, and frontoparietal control systems [42,46,50]. However, it remains unclear to what extent connectivity patterns that track one variety of arousal would also track others.

In the current work, we use predictive modeling of dynamic functional connectivity to situate varieties of arousal within a connectome-based neural reference space. Connectome-based predictive models (CPMs) predict individual traits or psychological states from whole-brain functional connectivity patterns [42,51–56]. We construct dynamic CPMs to predict continuous fluctuations in three varieties of arousal, following the taxonomy proposed by Satpute and colleagues [17]: affective arousal (subjective feelings of activation), autonomic arousal (peripheral physiological activity), and wakefulness arousal (vigilance and responsiveness). We situate different measures of arousal within this reference space in two complementary ways. First, we test whether models trained to predict one measure of arousal generalize to predict others. Successful generalization would indicate that different varieties of arousal share variance that can be predicted from dynamic functional connectivity. Second, we compare the predictive connections selected by each model to assess their similarity across measures of arousal. As the models are estimated in resting-state, movie-watching, and story-listening fMRI, both analyses also test whether generalization and overlap extend across intrinsic and stimulus-driven fluctuations in arousal. Convergence across rest, movies, and stories would indicate that the models are not merely tracking stimulus features associated with arousal or engagement with a stimulus.

Finally, we ask whether the arousal models extend beyond the waking states in which they were trained, and whether they recover a classic consequence of arousal for memory, namely that arousing events are better remembered than neutral ones [9–12]. To address the first question, we test their ability to predict stages of sleep associated with progressively lower levels of arousal. To address the second, we take advantage of the post-scan free recall in the movie datasets. We use each CPM to decode time-varying arousal while participants viewed the movies and test whether CPM-derived arousal estimates during a movie event predict how well the event is later remembered. Together, these analyses allow us to ask not only how different measures of arousal relate to one another within a connectome-based reference space, but also whether their shared variance, and the connectivity patterns that capture it, are behaviorally relevant.

## Results

We combined data from 98 participants drawn from five publicly available fMRI datasets [43,57–60] (**Figure 1A**). Across these datasets, participants contributed resting-state scans (N = 49), movie-watching scans (N = 29), or story-listening scans (N = 20). We regressed out white matter, cerebrospinal fluid, and global signal, as well as motion, from the BOLD time courses. For each participant, the fMRI data were parcellated into 268 cortical and subcortical regions of interest (ROIs) defined by the Shen atlas [61]. Dynamic functional connectivity (FC) was then estimated using a tapered sliding window of 30 seconds. Within each window, we computed the Fisher z-transformed Pearson correlation between the BOLD time series of every pair of ROIs.

**Figure 1.**
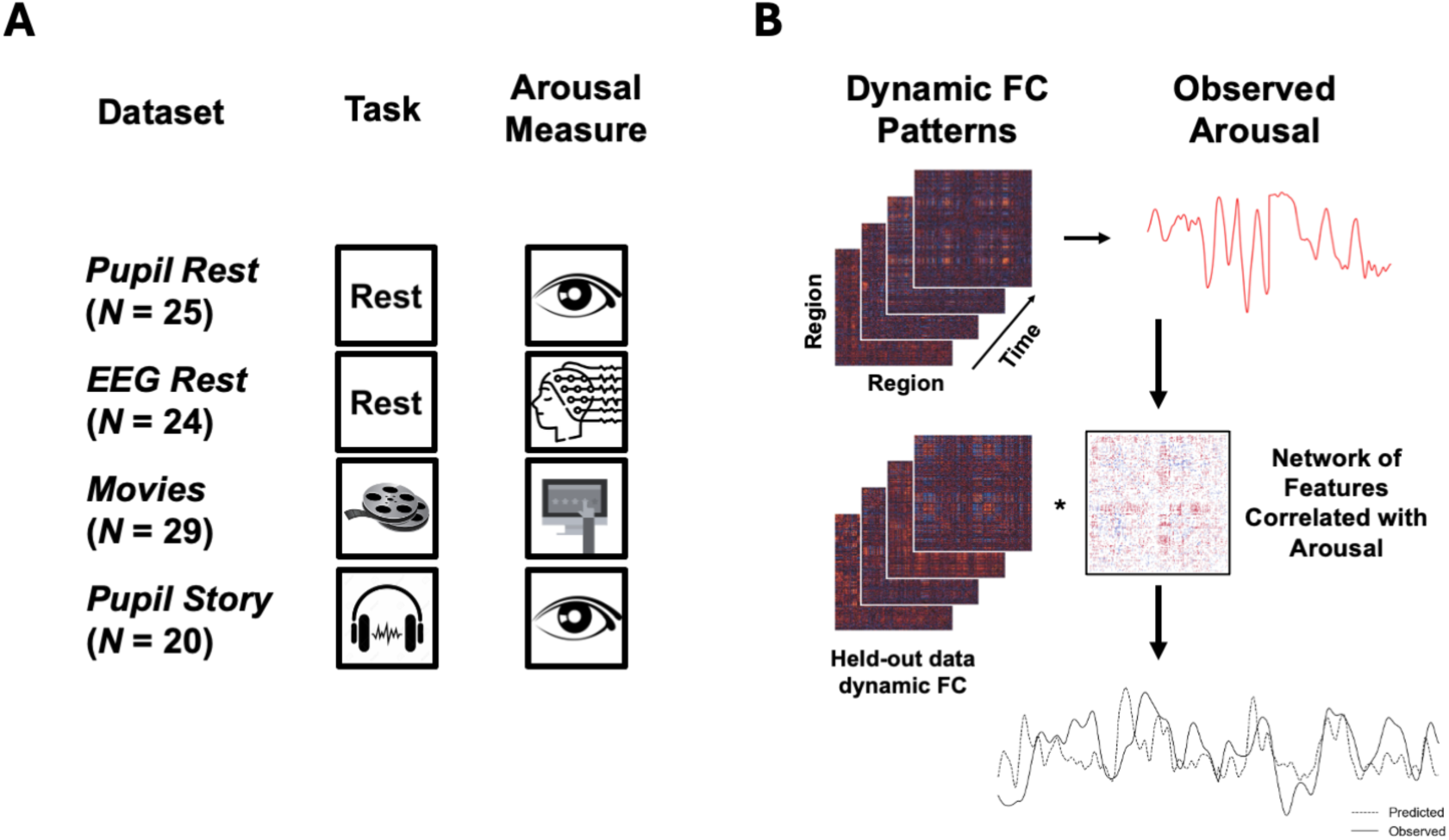
Overview of fMRI datasets and CPM pipeline. **(A)** The datasets included resting-state scans with concurrent measurements of pupil dilation (*Pupil Rest*) and EEG (*EEG Rest*), movie-watching scans (*Movies*) and story-listening scans (*Pupil Story*), with pupil dilation collected from a different group of participants listening to the same story. (**B**) Dynamic FC matrices were computed using a 30-second tapered sliding window. For each dataset, fluctuations in FC strength were correlated with the corresponding arousal time course to identify a network of connections reliably associated with arousal. We then calculated the element-wise product between this arousal network and held-out test data to generate predicted arousal time courses for each participant.

For each dataset, we then constructed an arousal network by correlating FC strength with a target arousal time course in a cross-validated manner, retaining only functional connections that showed a significant association at *p* < 0.05. Connections that were positively or negatively correlated with the target time course were assigned weights of +1 and -1, respectively, and all remaining connections were assigned a weight of 0. We then generated a predicted arousal time course by computing the element-wise product between the arousal network weights and the dynamic FC matrix at each time point of held-out test data (**Figure 1B**; see Methods).

## Dynamic CPMs generalize across pupil and EEG measures of arousal during wakeful rest

We first asked whether dynamic FC captures fluctuations in pupil dilation, a widely used measure of autonomic arousal [23]. Pupil size has also been found to be correlated with other autonomic measures such as heart rate and skin conductance [21]. Using an fMRI dataset with concurrent pupillometry during wakeful rest (*Pupil Rest*), we derived a pupil-rest CPM with a leave-one-participant-out cross-validation approach. The pupil dilation time courses were first convolved with a hemodynamic response function and smoothed using the same tapered sliding window used to estimate dynamic FC. In each cross-validation fold, we trained a model to predict fluctuations in pupil dilation from dynamic FC and tested it on the held-out participant. Across folds, average prediction accuracy was significantly higher than chance (mean *r* = 0.308, *p* < 0.001; **Figure 2**), with statistical significance assessed by comparing the average accuracy against a null distribution of 10,000 permutations in which the predicted pupil time courses were correlated with phase-randomized observed pupil time courses.

**Figure 2.**
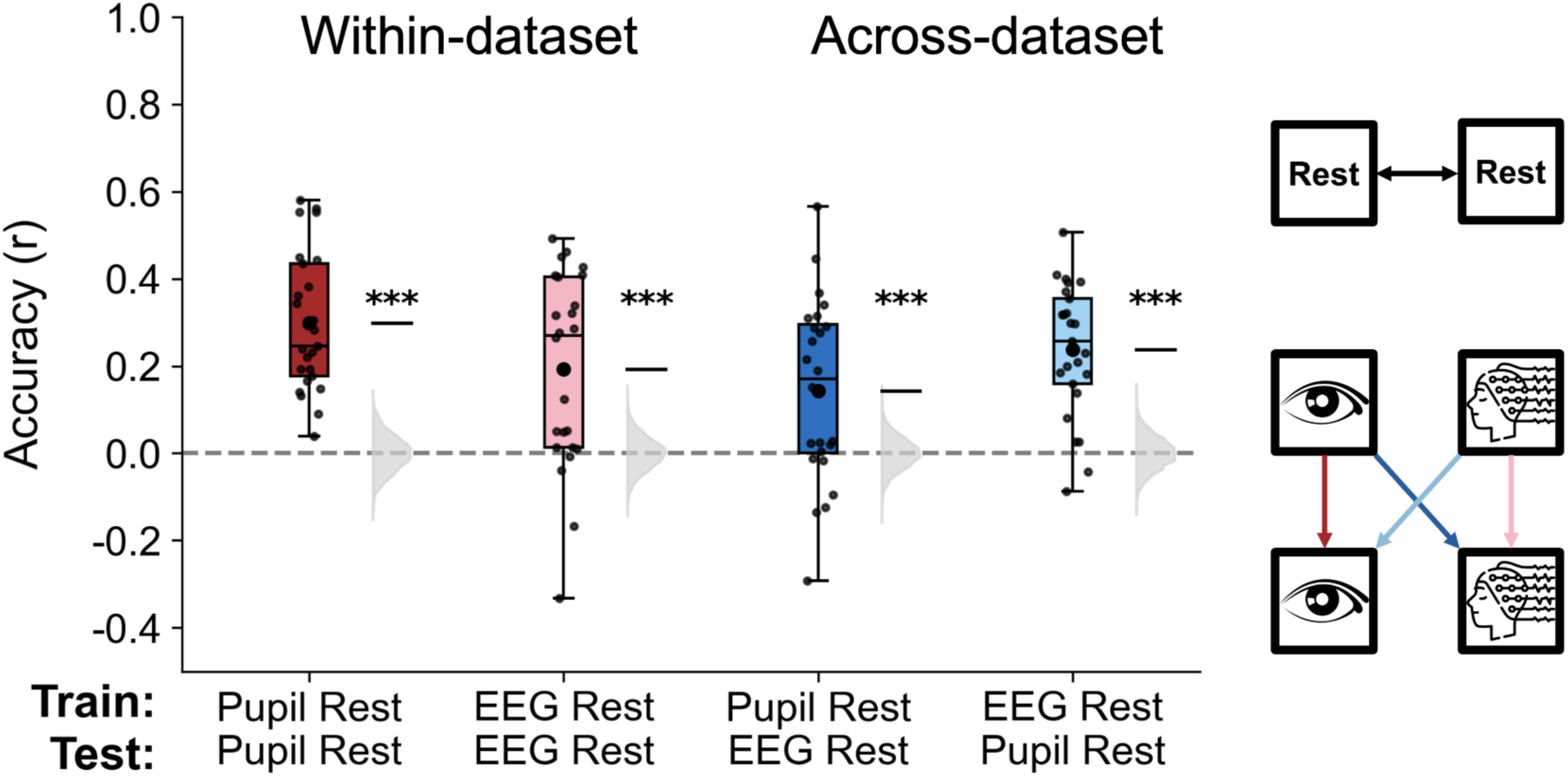
Dynamic CPMs generalize across pupil dilation and EEG spectral slope during wakeful rest. The y-axis displays model prediction accuracy, as measured by Pearson’s correlation between the model’s predicted time course and the observed arousal time course. Each datapoint within the box plot denotes a participant. The black horizontal lines show the average of the Fisher z-transformed *r*-values. The gray half-violin plots show the null distributions generated from correlating the predicted arousal time courses with phase-randomized observed arousal time courses. Red: Within-dataset prediction, Blue: Across-dataset prediction. \**p* < .05, ** *p* < .01, *** *p* < .001.

We next asked whether dynamic FC captures fluctuations in the spectral slope of the aperiodic component of the EEG power spectrum, an established measure of wakefulness arousal that tracks fluctuations in vigilance, transitions between wake and sleep, and changes in anesthesia depth [25,62–64]. Using a different fMRI dataset with concurrent EEG during wakeful rest (*EEG Rest*), we derived an EEG-rest CPM with the same leave-one-participant-out cross-validation procedure to predict fluctuations in EEG spectral slope. The EEG spectral slope time courses were convolved with a hemodynamic response function and smoothed using the same tapered sliding window used to estimate dynamic FC. Average prediction accuracy was significantly higher than chance (mean *r* = 0.201, *p* < 0.001; **Figure 2**).

Having found that dynamic FC can separately predict both pupil dilation and EEG spectral slope, we next asked whether a CPM trained on one arousal measure in one dataset generalizes to the other. Average prediction accuracy was significantly higher than chance both when training on pupil dilation and testing on EEG spectral slope (mean *r* = 0.150, *p* < 0.001) and when training on EEG spectral slope and testing on pupil dilation (mean *r* = 0.243, *p* < 0.001; **Figure 2**), indicating that FC patterns capture shared dynamics across pupil and EEG measures of arousal during wakeful rest.

We also assessed significance against a null distribution in which arousal time courses were shuffled across participants (see Methods). Within-dataset predictions of pupil dilation and EEG spectral slope remained significant against this null, as did cross-dataset predictions when training on pupil dilation and testing on EEG spectral slope and when training on EEG spectral slope and testing on pupil dilation (all *p*s < 0.001). These additional analyses suggest that above-chance prediction accuracy was not explained by temporal trends in arousal shared across participants (e.g., arousal systematically decreasing with increasing scan duration).

## Dynamic CPMs generalize across subjective affective arousal ratings and pupil dilation during movie-watching and story listening

Our previous work found that dynamic CPMs trained to predict subjective affective arousal ratings of one movie generalized to other movies [42]. Here, we extend this work to test if CPMs trained to predict subjective ratings of audiovisual movies would generalize to predict pupil dilation during story-listening, and vice versa. We combined two movie-watching datasets where participants watched audiovisual movies while undergoing fMRI (*Movies*). Each movie was segmented into discrete events defined by major shifts in the narrative (Movie 1: 48 events, average duration = 57.5 s; Movie 2: 68 events, average duration = 38.4 s). As a measure of affective arousal, we relied on subjective behavioral ratings for each event from a separate group of participants.

We then pooled participants across the two movie datasets and selected FCs that were consistently associated with arousal ratings across all participants. As the resulting combined movies CPM was trained on two different movies, its feature set was less likely to reflect idiosyncratic stimulus properties. We applied the combined movies CPM to an independent dataset in which participants listened to a 20-min spoken story while undergoing fMRI (*Pupil Story*). As this dataset did not include pupillometry data, we used pupil recordings from a separate group of participants listening to the same story and averaged pupil responses across participants to derive a normative time course of stimulus-evoked pupil dilation. Pupil fluctuations were previously shown to be correlated between participants in this dataset [46], indicating that the story evoked arousal in a reliable manner across participants.

The combined movies CPM significantly predicted this normative pupil time course when applied to the fMRI data during story listening (mean *r* = 0.136, *p* < 0.001; **Figure 3**). Similarly, a pupil-story CPM trained to predict the story-evoked pupil time course generalized to predict affective arousal ratings in the two movie datasets (mean *r* = 0.162, *p* < 0.001; **Figure 3**). Together, these results indicate that FC patterns that track affective arousal during movies generalize to predict autonomic arousal elicited by a story, and that FC patterns tracking story-evoked autonomic arousal generalize to predict affective arousal ratings during movies.

**Figure 3.**
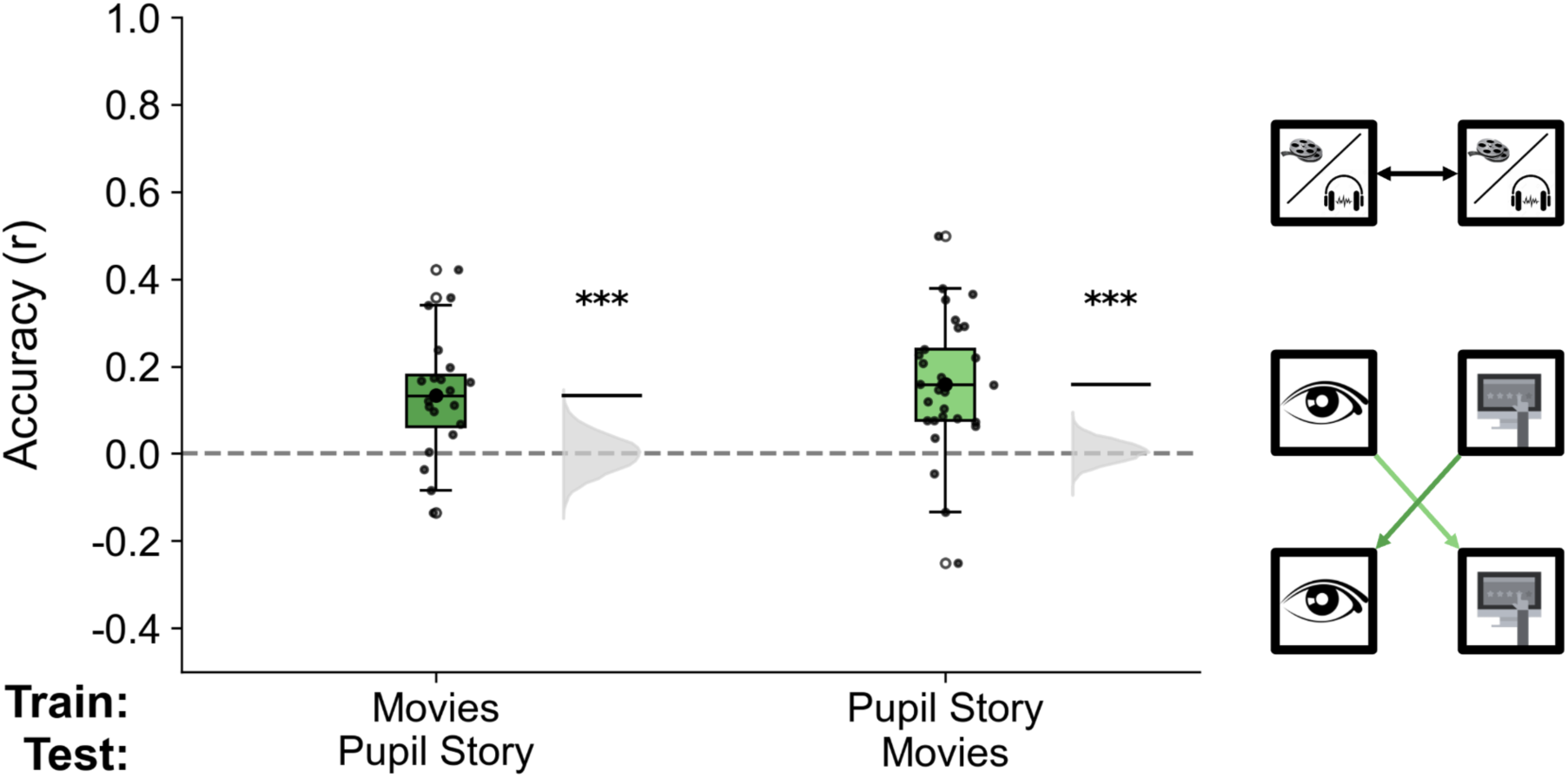
Dynamic CPMs generalize across subjective ratings and pupil dilation during movie-watching and story listening. The y-axis displays model prediction accuracy, as measured by Pearson’s correlation between the model’s predicted time course and the observed arousal time course. Each datapoint within the box plot denotes a participant. The black horizontal lines show the average of the Fisher z-transformed *r*-values. The gray half-violin plots show the null distributions generated from correlating the predicted arousal time courses with phase-randomized observed arousal time courses. \**p* < .05, ** *p* < .01, *** *p* < .001.

## Dynamic CPMs generalize across rest and narratives and across measures

We next asked whether CPMs also generalize between wakeful rest and narratives, which differ in whether arousal fluctuates spontaneously or in response to an external stimulus. We first tested whether CPMs trained to predict arousal fluctuations during wakeful rest generalize to stimulus-evoked arousal during narratives. Both models that were trained on resting-state data generalized to predict subjective ratings during movie-watching (pupil-rest CPM: mean *r* = 0.151, *p* < 0.001; EEG-rest CPM: mean *r* = 0.161, *p* < 0.001; **Figure 4A**) and pupil dilation during story-listening (pupil-rest CPM: mean *r* = 0.121, *p* = 0.001; EEG-rest CPM: mean *r* = 0.169, *p* < 0.001; **Figure 4A**).

**Figure 4.**
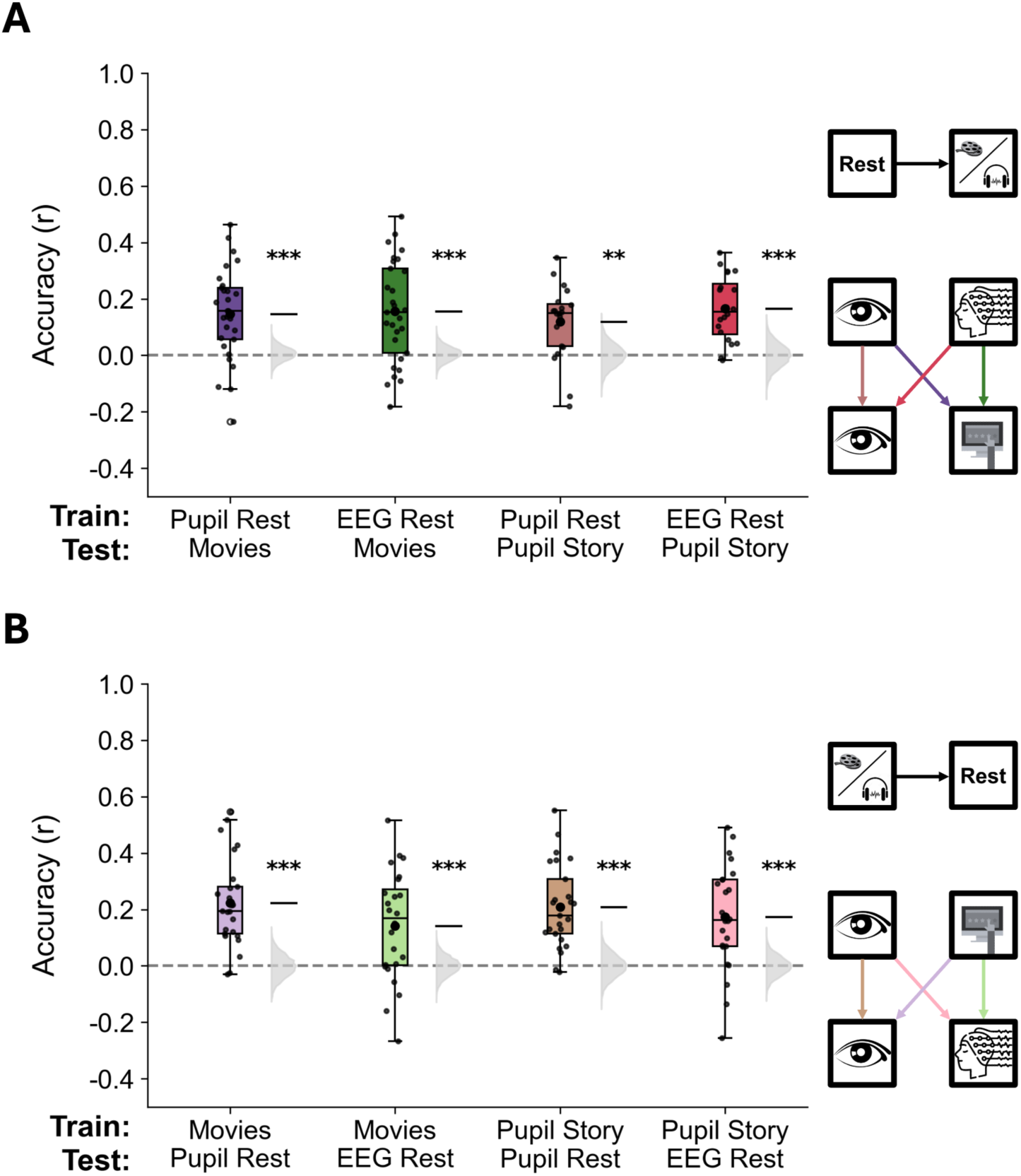
Dynamic CPMs generalize across rest and narratives and across measures. **(A)** CPMs trained on spontaneous arousal fluctuations during rest significantly predicted subjective ratings during movie-watching and pupil dilation during story listening. **(B)** CPMs trained on subjective ratings during movie-watching and pupil dilation during story listening significantly predicted spontaneous arousal fluctuations during rest. In each panel, the y-axis displays the prediction accuracy, as measured by Pearson’s correlation between the model’s predicted time course and the observed arousal time course. Each datapoint within the box plot denotes a participant. The black horizontal lines show the average of the Fisher z-transformed r-values. The gray half-violin plots show the null distributions generated from correlating the predicted arousal time courses with phase-randomized observed arousal time courses. \**p* < .05, ** *p* < .01, *** *p* < .001.

We next tested whether CPMs trained on stimulus-evoked arousal during narratives generalize to spontaneous arousal fluctuations during wakeful rest. Indeed, CPMs trained on subjective ratings during movie-watching and pupil dilation during story-listening generalized to predict pupil dilation (movies CPM: mean *r* = 0.230, *p* < 0.001; pupil-story CPM: mean *r* = 0.215, *p* < 0.001) and EEG spectral slope (movies CPM: mean *r* = 0.148, *p* < 0.001; pupil-story CPM: mean *r* = 0.179, *p* < 0.001; **Figure 4B**). These predictions also remained significant against null distributions in which arousal time courses of pupil-rest and EEG-rest were shuffled across participants (all *p*s < 0.001).

To contextualize these cross-dataset accuracies, we compared them with within-dataset accuracy. For each model, we computed a generalization ratio, defined as the cross-dataset accuracy divided by the within-dataset accuracy for the test dataset (**Table S1**). A ratio of one would indicate that a model trained on a different measure and context predicts the test measure as well as a model trained on the measure itself. When testing on resting-state datasets, generalization ratios ranged from 0.70 to 0.79 for pupil dilation and from 0.74 to 0.89 for EEG spectral slope. Ratios were lower when testing on narrative datasets, ranging from 0.44 to 0.48 for movie ratings and from 0.48 to 0.68 for story-evoked pupil dilation. The lower ratios likely reflect within-dataset models in the narrative datasets capturing both arousal and stimulus features that covary with arousal in each narrative [42,53]. Models trained on other datasets would not capture stimulus-specific features, resulting in lower cross-dataset prediction accuracies.

We next assessed whether these results depended on the choice of parcellation, sliding-window length, or global signal regression. We first repeated the analysis using the Schaefer 200-region cortical parcellation [65] combined with 8 Brainnetome subcortical regions [66] (bilateral amygdala, hippocampus, basal ganglia, and thalamus). Within-dataset and across-dataset prediction accuracies were significantly above chance for all train and test combinations (**Table S1**). We then reran all cross-dataset predictions using shorter (16-second) and longer (44-second) sliding windows, and all cross-dataset prediction accuracies remained significantly above chance (all *p*s < 0.01; **Figures S1 and S2; Table S2**). Finally, we repeated the analysis without global signal regression. Within-dataset accuracies remained significant for all datasets, and 9 of 12 cross-dataset predictions remained significant, although accuracies were generally lower (**Table S3**). The 3 exceptions involved the Pupil Story dataset (pupil-rest CPM to Pupil Story, EEG-rest CPM to Pupil Story, and pupil-story CPM to Pupil Rest). With the Schaefer-Brainnetome parcellation, 11 of 12 cross-dataset predictions remained significant, with the pupil-rest CPM to Pupil Story the only exception (Table S3).

## Shared predictive connections and similarity of edge-level associations across models

Generalization across contexts and measures provides evidence that connectivity patterns track shared arousal dynamics across spontaneous and stimulus-evoked states, as well as across affective, autonomic, and wakefulness measures. To characterize these connectivity patterns more directly, we examined the predictive connections (i.e., edges) selected by each model.

We first computed the proportion of two-way, three-way, and four-way overlap among the predictive edges selected in the pupil-rest, EEG-rest, pupil-story, and movies CPMs. Edges were considered overlapping if they correlated with arousal in the same direction across models. To determine whether the observed overlaps exceeded what would be expected by chance, we used a permutation procedure where, for each model, we randomly selected the same number of positive and negative edges that were selected in the corresponding model and recomputed overlap to generate a null distribution of overlap proportions [67]. The proportion of overlapping edges was significant for every possible combination of models (all *p*s < 0.001; **Figure 5**), and this was also the case with the Schaefer-Brainnetome parcellation (all ps < 0.001), indicating that the CPMs drew on partially shared sets of functional connections.

**Figure 5.**
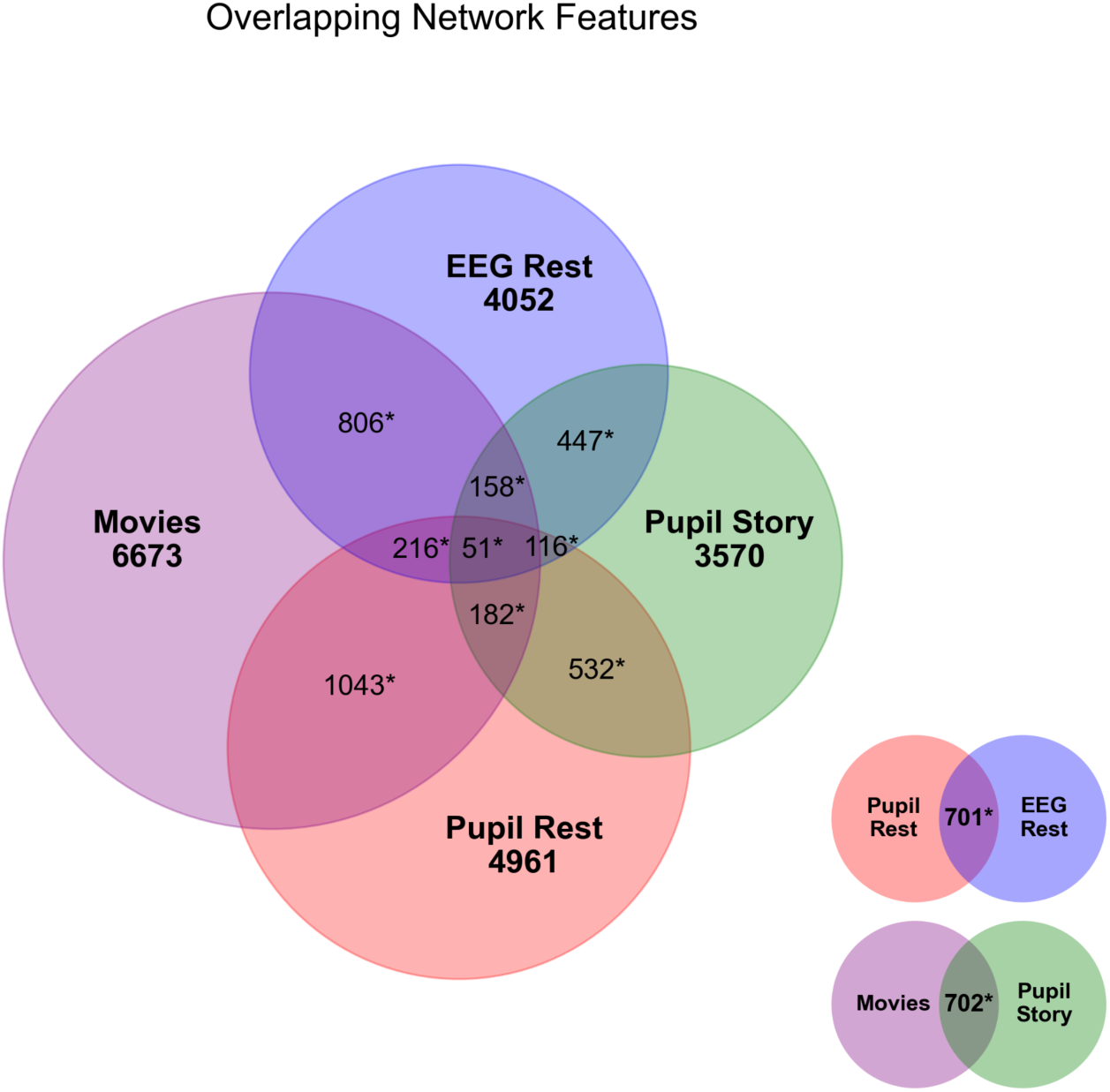
Number of unique and overlapping FCs among arousal networks. Features were considered overlapping if they were significantly correlated with arousal in the same direction between networks. * indicates a significant proportion of overlap at *p* < 0.001.

There were 51 predictive edges that were common to all four models (**Figure 6A, Table S4**; see **Figure S3, Table S5** for edges common to all four models when using the Schaefer-Brainnetome parcellation). A predictive model restricted to these edges significantly predicted arousal fluctuations in every dataset (**Figure S4A**). As the four-way overlap includes edges selected using each test dataset, we also tested a stricter version of this analysis, in which a model restricted to the edges shared by the other three models was used to predict arousal in the held-out dataset. These models also significantly predicted arousal in every held-out dataset (**Figure S4B**).

**Figure 6.**
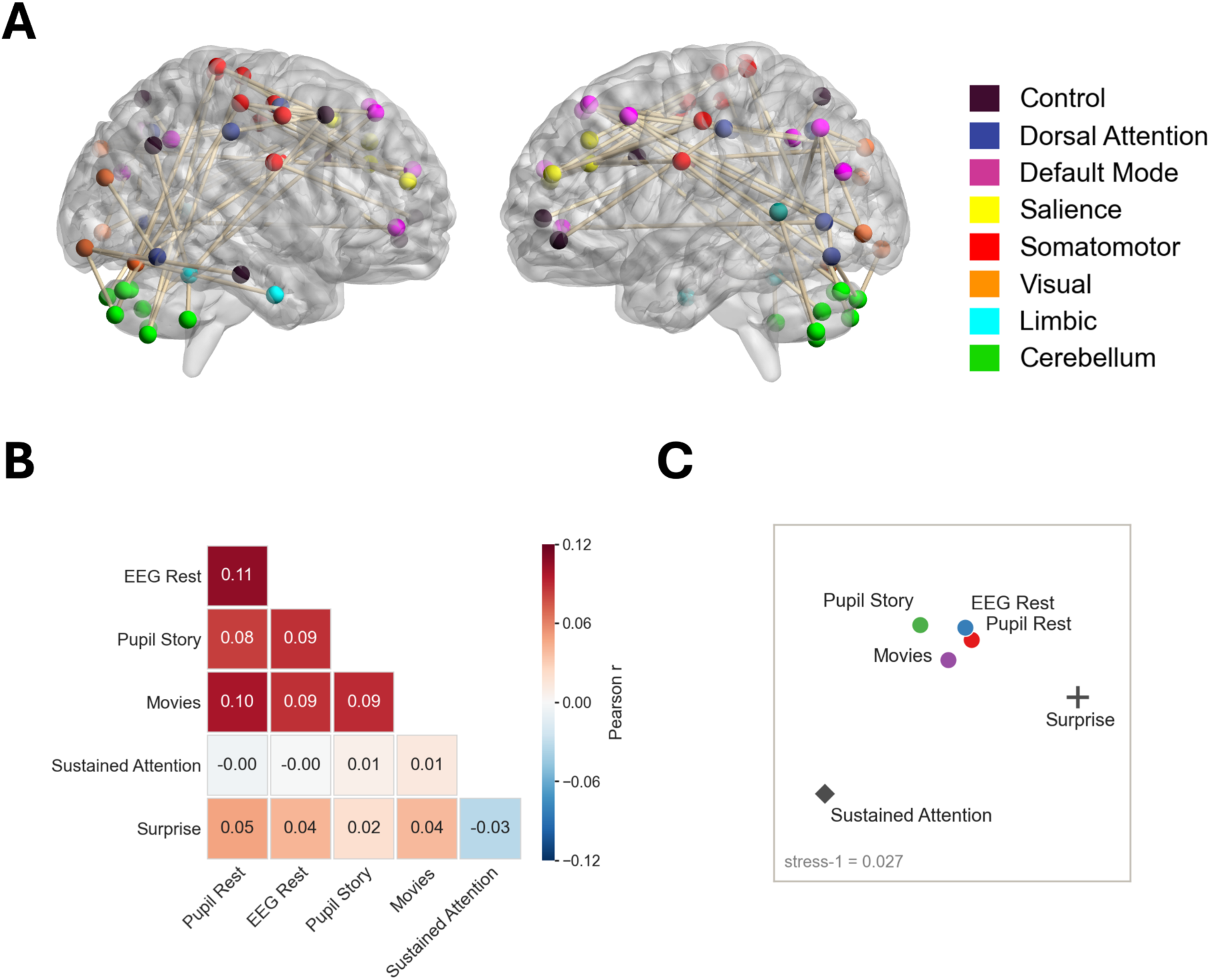
Shared predictive connections and similarity of edge-level associations across models. **(A)** Glass brain displaying 51 functional connections that overlap across all four CPMs (Shen 268 atlas, feature selection at p < 0.05). Nodes indicate brain regions, with colors denoting functional networks. **(B)** Pearson correlations between the edge-level association maps of the four arousal CPMs and of published connectivity models of sustained attention [51,68] and surprise [54]**. (C)** Two-dimensional embedding of the similarities between the feature maps of the models obtained with multidimensional scaling, with distance defined as one minus Pearson correlation. A stress-1 value of 0.027 indicates that the embedding closely preserves the pairwise distances. Statistical comparisons of the similarities are shown in Figure S6.

All common predictive edges using the Shen atlas were positive and connected 53 unique nodes distributed across multiple canonical functional networks. We compared the proportion of edges within and between networks to a null distribution generated by shuffling the edges while preserving the number of significant ROIs. Connections between the salience and somatomotor networks (10 edges, *q* = 0.004) and between the somatomotor and cerebellar networks (8 edges, *q* = 0.011) were overrepresented relative to chance. However, these network-level patterns were not consistent when using the Schaefer-Brainnetome atlas, across feature-selection thresholds, and without global signal regression (**Figure S5**). None of the common predictive edges involved a subcortical region in either parcellation. To further assess the contribution of subcortical regions, we examined the edges selected by each individual model and those shared by any three of the four models. As ROIs in the Shen parcellation are not anatomically labeled, we focused this analysis on the Schaefer-Brainnetome parcellation. Subcortical edges were present in all four individual models and in every three-way overlap, and each of the eight subcortical regions appeared in at least one three-way overlap (**Table S6**).

To quantify the similarity in predictive connectivity patterns for each pair of arousal models, we computed the Pearson correlation between the edge-level features of each model (**Figure 6B**). For each measure, we represented the associations between every edge and arousal as a signed map (thresholded at *p* < 0.05), assigning +1 to edges positively associated with arousal, -1 to edges negatively associated with arousal, and 0 to all other edges. To place these similarities in context, we also computed the correlation with edges in connectome-based predictive models that have been previously shown to predict continuous fluctuations in two psychological states, sustained attention [51,68] and surprise [54]. Both the sustained attention and surprise models were originally defined in the Shen atlas, and, like the arousal CPMs in our main analyses, were trained on data from which the global signal had been regressed, and consist of positive and negative networks of edges selected by their correlation with the target state.

We visualized the relationships among these models by embedding the pairwise similarities in two dimensions using multidimensional scaling, with distance defined as one minus the Pearson correlation (**Figure 6C**). The four arousal models were positioned close to one another, with surprise at an intermediate distance and sustained attention farther away. To test this pattern statistically, we compared the mean similarity among the arousal models with their mean similarity to each comparison model, using a bootstrap over participants to compute *p*-values. The arousal models were more similar to one another (mean *r* = 0.092) than to the surprise model (mean *r* = 0.037; difference = 0.056, *p* < 0.001) or the sustained attention model (mean *r* = 0.004; difference = 0.089, *p* < 0.001), and more similar to the surprise model than to the sustained attention model (difference = 0.033, *p* = 0.003; **Figure S6**). The same pattern was observed at different feature-selection thresholds, with unthresholded arousal feature maps, after projecting the comparison models to the Schaefer-Brainnetome parcellation scheme, without global signal regression, and with different sliding-window lengths (**Figure S6; Figure S7)**.

## Dynamic CPMs generalize to graded arousal across sleep stages

We next asked whether the predictive functional connections across arousal varieties extend to the graded fluctuations in arousal that accompany transitions into sleep. To test this, we applied the arousal CPMs to an *EEG Sleep* dataset where participants underwent simultaneous EEG and fMRI during 15-minute sleep sessions [60]. Each 30-second epoch had been manually scored as Wake, NREM1, NREM2, or NREM3, reflecting progressively deeper sleep and decreasing arousal. All four models significantly predicted sleep stage from dynamic FC (pupil-rest CPM: mean *r* = 0.107, *p* < 0.001; EEG-rest CPM: mean *r* = 0.091, *p* < 0.001; pupil-story CPM: mean *r* = 0.109, *p* < 0.001; movies CPM: mean *r* = 0.078, *p* < 0.001; **Figure 7)**. A model restricted to the edges shared across all four CPMs (the four-way network) also significantly predicted sleep stage (mean *r* = 0.070, *p* = 0.001). For the EEG-rest CPM, which was trained on the same participants whose sleep data were tested, predictions were obtained using leave-one-participant-out cross-validation, where the EEG-rest CPM was trained on all participants except the test participant.

**Figure 7.**
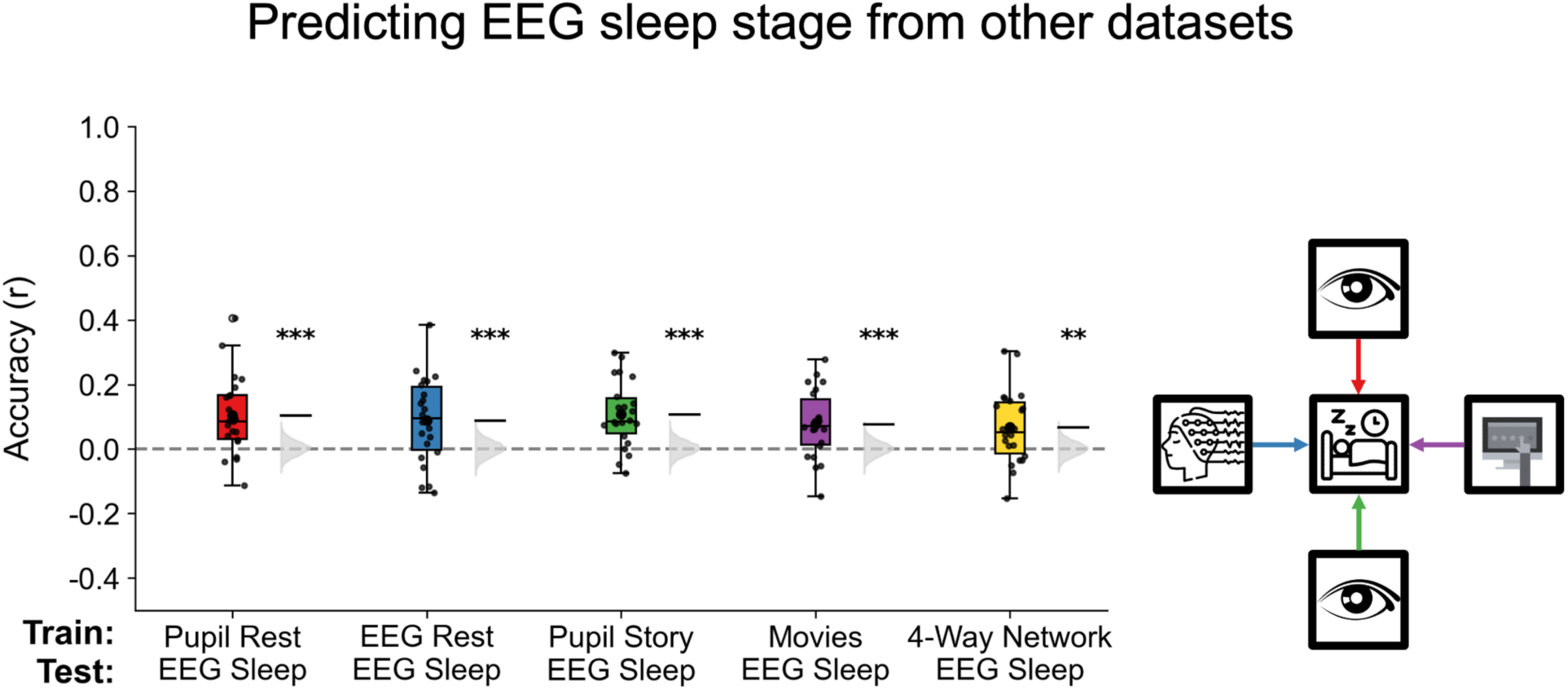
Dynamic CPMs generalize to graded arousal across sleep stages. Each CPM was applied to predict manually scored sleep stages in the *EEG Slee*p dataset. The y-axis displays model prediction accuracy, as measured by Pearson’s correlation between the model’s predicted arousal time courses and the manually scored sleep stage time courses. Each datapoint within the box plot denotes a participant. The black horizontal lines show the average of the Fisher z-transformed *r*-values. The gray half-violin plots show the null distributions generated from correlating the predicted sleep stage time courses with phase-randomized sleep-stage time courses. \**p* < .05, ** *p* < .01, *** *p* < .001.

## Decoded arousal during movie-watching predicts subsequent recall fidelity

Having established that the arousal CPMs generalize across measures and contexts, we next asked whether the decoded arousal dynamics were behaviorally meaningful. We had selected these two movie-watching datasets because they included post-scan free recall, in which participants verbally recounted what they remembered about the movie. This allowed us to test whether CPM-decoded arousal during movie-viewing reproduces well-established arousal-related memory enhancement effects.

We used Google’s Universal Sentence Encoder (USE) [69] to convert detailed annotations of each movie event and participants’ recall transcripts into numerical vectors that reflect the semantic meaning of the text. Following previous work [46,70,71], we quantified recall fidelity as the cosine similarity between each event’s annotation vector and the vector of a participant’s recall of the corresponding event. We then used Bayesian multilevel models to predict the recall fidelity of an event from decoded arousal while viewing the event, as decoded by each CPM.

For each CPM, higher decoded arousal while viewing a movie event was associated with higher subsequent recall fidelity, consistent with the well-established association between arousal and memory (pupil-rest CPM: b = 0.064, p(b > 0) = 0.992, 95% HDI [0.016, 0.118], WAIC = 4595; EEG-rest CPM: b = 0.083, p(b > 0) = 0.999, 95% HDI [0.033, 0.137], WAIC = 4590; pupil-story CPM: b = 0.054, p(b > 0) = 0.984, 95% HDI [0.003, 0.102], WAIC = 4596; movies CPM: b = 0.080, p(b > 0) = 0.999, 95% HDI [0.030, 0.128], WAIC = 4592; WAIC_null_ = 4600; **Figure 8**). For the movies CPM, decoded arousal was obtained using leave-one-participant-out cross-validation, in which the model was trained on all participants except the one whose recall was being predicted. Decoded arousal from the four-way network also predicted recall fidelity (b = 0.058, p(b > 0) = 0.989, 95% HDI [0.010, 0.107], WAIC = 4595), indicating that the shared signal captured by these models is predictive of subsequent memory.

**Figure 8.**
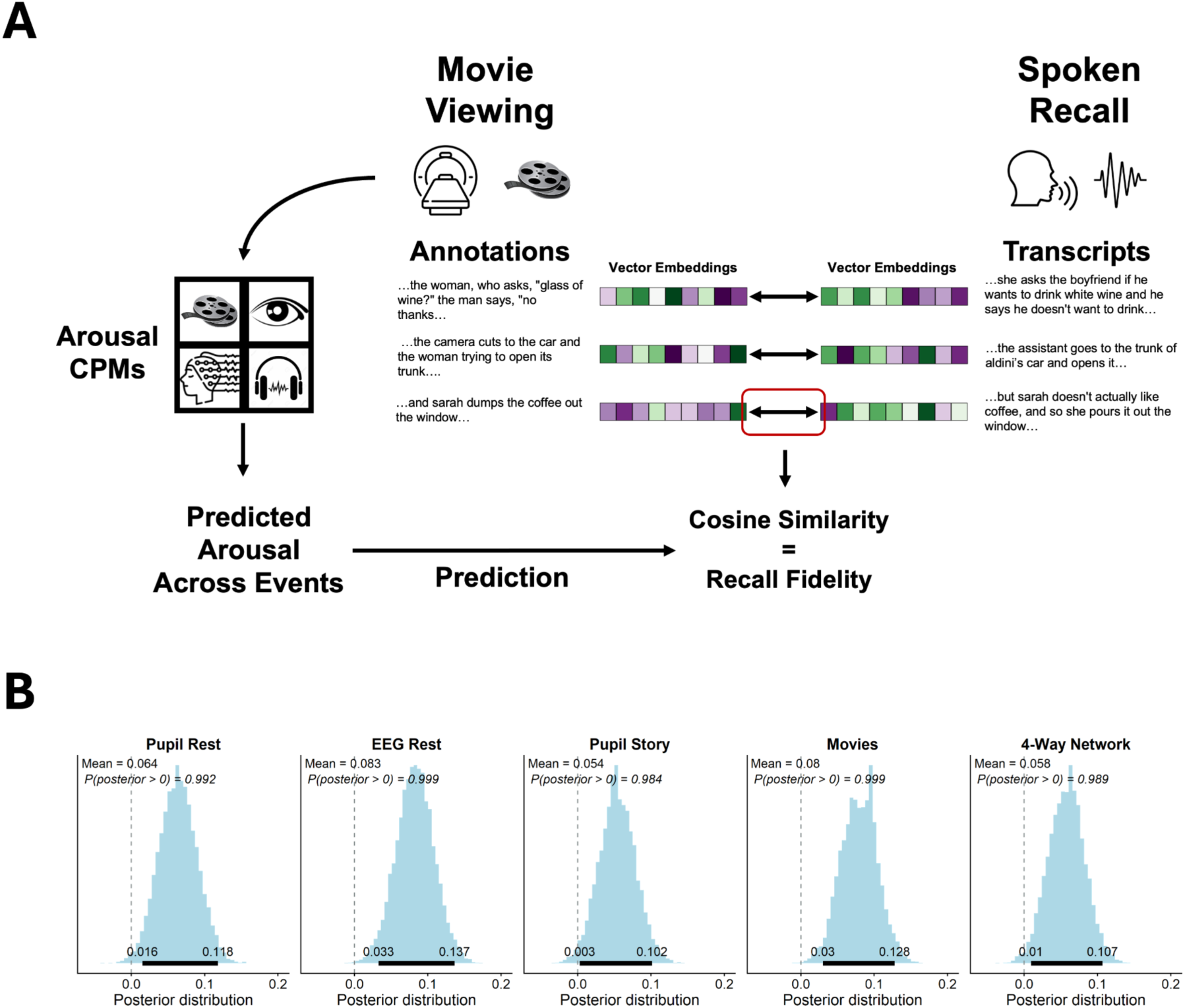
Predicting subsequent memory from decoded arousal during movie-watching. **(A)** Movie annotations and participants’ recall transcripts were converted into sentence embeddings. Recall fidelity was computed as the cosine similarity between the sentence embeddings of a recalled event and the matching movie annotation. For each event, we obtained five separate measures of decoded arousal by applying the movies, pupil-rest, pupil-story, and EEG-rest CPMs, and the four-way network to the movie-watching fMRI data. Each decoded arousal measure was used to predict recall fidelity of the matching event using a Bayesian multilevel model. **(B)** Posterior distributions of the effect of each decoded arousal measure on recall fidelity. Thick solid lines indicate the 95% highest density interval.

## Discussion

Across wakeful rest, movie-watching, and story-listening, fluctuations in measures of affective, autonomic, and wakefulness arousal were significantly predicted from patterns of dynamic connectivity. Predictive models trained on one measure and task context generalized to the others, and the connections selected by each model overlapped more than expected by chance. The models also predicted graded changes in arousal across sleep stages, and arousal decoded during movie-watching predicted how well a movie scene was later remembered. By situating different measures of arousal in a connectome-based neural reference space, we show that the relationship among them can be characterized empirically, revealing partial convergence that extends across spontaneous and stimulus-driven fluctuations.

A strength of the current approach is that it demonstrates generalization across both spontaneous fluctuations in arousal during wakeful rest and arousal elicited by external narratives. Stimulus features often covary with experienced arousal [72–74], raising the possibility that decoders trained on movies or stories track properties of the stimulus rather than arousal. Consistent with this concern, human arousal ratings can be predicted from stimulus features extracted by computer vision models that do not have bodily states or subjective experience [75], indicating that stimulus structure alone can account for variance in normative arousal judgments. At the same time, arousal decoded during rest and sleep is also susceptible to alternative explanations, including head motion, scanner drift, or respiration-related coupling between physiological signals and the BOLD response [76–78]. Generalization across stimulus-viewing, rest, and sleep therefore provides greater confidence that the decoded signal tracks arousal-related brain states, rather than stimulus-related confounds or non-neuronal contributions.

The cross-dataset prediction accuracies we observed are comparable to those reported in prior work using dynamic CPMs to predict fluctuations in psychological states across datasets [42,53,54,56,79], and are bounded by the reliability of both the arousal measures and fMRI. Notably, models trained on other measures and contexts achieved a meaningful fraction of the accuracy of models trained on the test measure itself, ranging from roughly half to most of it depending on the test dataset. Nevertheless, the prediction accuracies were modest in absolute terms, and each measure was still best predicted by a model trained on that measure within the same dataset. The shared signal we identify is therefore consistent across measures but limited in magnitude, and our results should not be taken to imply that different varieties of arousal are neurally identical or reducible to a single unitary process. We interpret this pattern of modest but consistent generalization as partial convergence across measures of arousal, in which different measures track partly overlapping dynamics in brain connectivity.

The partial nature of this convergence is consistent with long-standing evidence that measures of arousal can sometimes diverge. Even when recorded concurrently in the same individuals, moment-to-moment ratings of subjective experience are only modestly associated with cardiovascular and electrodermal responses [28,29], and different autonomic measures can change in opposite directions [23,31]. Our results suggest that a similar pattern of partial correspondence extends to the patterns of brain connectivity associated with these measures. This partial correspondence was not observed in a recent study by Zhang and colleagues [39], which derived an activation-based multivariate signature of affective arousal and found that it did not generalize to predict autonomic activation or states of wakefulness. One possibility is that regional activity patterns better discriminate between varieties of arousal, whereas brain-wide changes in connectivity that accompany shifts in arousal state are more likely to be shared. Such changes in connectivity patterns need not reflect interactions between specific regions. Recent work suggests that arousal-related changes in functional connectivity may arise from a common arousal-related process that modulates activity across many regions together [47,49]. Dynamic FC, which captures coordinated fluctuations between brain regions, would be more sensitive to these shared changes.

In our main analysis, 51 functional edges were shared across all four models, and both network pairs that were overrepresented relative to chance included the somatomotor network. This is consistent with prior resting-state fMRI studies showing that the somatomotor cortex is among the regions most sensitive to arousal [80–82]. Stimulant medication, which increases arousal, has also been associated with increased connectivity between somatomotor and salience regions [83]. We note, however, that the distribution of edges across networks varied across analytical choices, in line with a recent study showing that univariate feature selection in brain-based predictive models can yield different sets of features with similar predictive performance [84]. The shared edges are therefore best understood as one snapshot of a distributed pattern, in which different edges emerge as top features depending on analytical choices, rather than as a unique neural substrate of arousal.

The same caution applies to the absence of particular regions from the shared edges. Despite past studies implicating subcortical regions across varieties of arousal [26,80,85,86], none of the edges shared across all four models involved a subcortical region. Subcortical edges, however, were present in all four individual models and in every three-way overlap, with each of the eight subcortical regions appearing in at least one three-way overlap. This suggests that subcortical edges were excluded by the stringent requirement that an edge be selected by all four models. Our parcellation schemes may have represented the subcortex too coarsely, combining nuclei with distinct functions [87]. Furthermore, many subcortical regions have a lower signal-to-noise ratio and are more susceptible to imaging artifacts [88–90]. These factors could reduce how consistently subcortical edges are selected across models. Their absence from the shared edges should therefore not be taken as evidence that subcortical regions are not associated with arousal.

When we compared the feature maps across models, the arousal models were more similar to one another than to models of sustained attention [51] and surprise [54], and more similar to surprise than to sustained attention. The intermediate position of surprise is consistent with evidence that surprising events evoke phasic increases in arousal [91,92], and models of surprise and arousal may therefore capture partly overlapping fluctuations in brain connectivity. In contrast, sustained attention may not be linearly related to arousal, as lapses are likely when arousal is either too high or too low [93–95]. Thus, fluctuations in arousal and sustained attention may be associated with distinct patterns of brain connectivity. One caveat in interpreting these differences is that we used published models of sustained attention and surprise rather than re-deriving them with our analytic pipeline. While the pattern of results was robust across parcellations, feature-selection thresholds, and preprocessing choices, differences in how the models were originally derived could still contribute to differences in similarity.

In particular, the sustained attention model was trained to predict individual differences in sustained attention from static functional connectivity [51]. It has since been shown to predict within-person changes in attention during an fMRI session [68], suggesting that its features covary dynamically with sustained attention. However, training the model on individual differences may have prioritized features that vary more across individuals than within individuals over time. Future work could compare arousal with models of other states built using the same analytic pipeline, which would help separate differences between psychological states from differences in how the models were derived. Extending this comparison to a broader range of psychological states would also provide a more complete picture of how arousal relates to other states within the connectome-based reference space.

Our findings complement two recent studies investigating the relationship between dynamic FC, arousal, and memory. Huang and colleagues [50] used a similar dynamic CPM approach to identify separate connectivity networks that predict arousal ratings of images and subsequent memory of those images, finding that pharmacologically inducing arousal by administering cortisol increased the overlap between the functional connections predictive of arousal and those predictive of memory. In a second study [46], our research group applied graph-theoretic analyses to the same movie-watching and story-listening datasets used in the current study, showing that high-arousal moments are associated with increased brain-wide integration across functional brain networks, which in turn predicted better subsequent memory for those moments. The current findings extend this work by showing that connectivity patterns associated with spontaneous fluctuations in autonomic and wakefulness arousal during rest, as well as the connections shared across different measures of arousal, also predict which moments are later remembered.

One limitation of the current study is that each variety of arousal was indexed by a single measure, and our results for each measure should not be taken as representative of that variety as a whole. Future work could situate multiple measures of the same variety of arousal in the same reference space, or examine the same measure across different contexts. As the reference space does not assume that measures converge, such work could identify where measures diverge as well as where they converge. A second limitation concerns the global fMRI signal, which has been previously shown to covary with physiological measures of arousal [20]. We regressed it out in our main analyses to ask whether arousal could be predicted from patterns of connectivity beyond the global signal, but doing so may also remove arousal-related variance along with non-neuronal noise. Most cross-dataset predictions remained significant without global signal regression, although the distribution of shared edges across networks differed. The choice of preprocessing should therefore be considered when comparing our findings with those of other studies.

In sum, we provide a connectome-based account of how multiple operationalizations of arousal relate to one another in the brain. Applying dynamic CPMs across wakeful rest and naturalistic narratives, we show that affective, autonomic, and wakefulness arousal can be predicted from time-varying functional connectivity, that these models generalize across measures and contexts, and that their predictions draw on partially overlapping sets of functional connections. These results suggest that different measures of arousal partially converge within a connectome-based reference space. More broadly, this framework offers a path toward moving beyond debates about whether arousal is “unitary” or “heterogeneous” by providing quantitative tools to map how arousal varieties converge and diverge depending on task demands, stimulus modality, and individual differences. Extending this approach to clinical populations and pharmacological manipulations may help identify conditions under which arousal-related network states become amplified or suppressed [83,96], clarifying how arousal relates to cognition.

## Materials and Methods

### Ethics Statement

Data analysis was approved by the Institutional Review Board of the University of Chicago (IRB22-0002). All data were anonymized by the original authors of each dataset. Data collection procedures were approved by the Institutional Review Board of the respective institutions. Written informed consent was obtained from all participants.

### fMRI Datasets

Five fMRI datasets from OpenNeuro were analyzed in this study. Across all datasets, participants with either more than 3 mm of translation or more than 3 degrees of rotation for any fMRI run were excluded from the analyses.

#### Pupil Rest

The Lee et al. (2022) Yale Resting-State Pupillometry/fMRI dataset [43] includes two runs of simultaneous fMRI and pupil dilation data collected during wakeful rest from 27 participants (16 female, 11 male; mean age = 26.5 years). Following the original authors, we discarded the first 10 s of each run, yielding 13 min 20 s of usable data (TR = 1 s, TE = 30 ms, flip angle = 55°, voxel size = 2x2x2 mm^3^). Two participants were excluded based on our motion exclusion criteria, yielding a final sample of 25 participants.

#### EEG Rest

The Gu et al. (2023) simultaneous EEG-fMRI dataset [60] includes two resting-state runs with concurrent EEG recordings collected during wakeful rest, yielding 20 min of usable fMRI data (TR = 2.1 s, TE = 25 ms, flip angle = 90°, voxel size = 3x3x4 mm^3^) from 33 participants (16 female, 17 male; mean age = 22.1 years). Seven participants were excluded due to clock synchronization and BESA-related file errors in the EEG data, and two additional participants were excluded based on our motion criteria. This resulted in a final sample of 24 participants.

#### Pupil Story

The Finn et al. (2018) *Paranoia* dataset [59] includes fMRI data acquired while 22 participants (11 female, 11 male; mean age = 27.0 years) listened to a ∼20-min audio story (TR = 1 s, TE = 30 ms, flip angle = 60°, voxel size = 2x2x2 mm^3^), presented across three fMRI runs. Based on our motion exclusion criteria, we included 20 participants. As the original *Paranoia* dataset did not include pupil recordings, we used a behavioral dataset collected by Park et al. (2025) [46], where an independent sample of 35 participants listened to the same audio story while their pupil size was recorded. Following the original authors, we excluded three participants due to equipment failure and five participants for noisy pupil data (i.e., more than a quarter of the data points deviated from the median pupil size by 2 standard deviations).

#### Movies

Movie-watching data were drawn from two publicly available datasets: the Chen et al. (2017) *Sherlock* dataset (Movie 1, TR = 1.5 s, TE = 28 ms, flip angle = 64°, voxel size = 3x3x4 mm^3^) [57], in which 17 participants (7 female, 10 male; mean age = 20.8 years) watched one 50-min TV episode, and the Lee et al. (2022) *Film Festival* dataset (Movie 2, TR = 1.5 s, TE = 39 ms, flip angle = 50°, voxel size = 2x2x2 mm^3^) [58], in which 21 participants (12 female, 9 male; mean age = 26.6 years) watched 10 short films with a collective duration of 50 minutes. Following the original authors, the cartoon introduction at the beginning of each run was discarded. The time courses were shifted by 4.5 s to account for hemodynamic lag. In *Sherlock*, one participant was excluded due to missing data, one due to preprocessing errors, and one due to our motion criteria. In *Film Festival*, we excluded three participants according to our motion criteria and three additional participants who were excluded by the original authors. In total, we included 29 of 38 participants across the two datasets.

We obtained event segmentations of *Sherlock* and *Film Festival* from [57] and [46], respectively, where the movie was divided into events defined by major shifts in the narrative, including changes in topic, location, time, and characters (*Sherlock:* 48 events, average length = 57.5 s, SD = 41.8 s; *Film Festival*: 68 events, average length = 38.4 s, SD = 18.2 s). We used behavioral arousal ratings collected in [46], where a separate group of 30 participants rated the arousal of each event on a scale of 1 to 5. The ratings were z-scored within each participant, and averaged across the group to obtain the arousal rating for each event.

### MRI preprocessing

MRI data were processed using fMRIPrep [97]. For anatomical data, the T1w image was corrected for intensity non-uniformity and skull-stripped. Brain tissue segmentation of cerebrospinal fluid, white matter, and gray matter was performed on the resulting image. Volume-based spatial normalization to the MNI152NLin6Asym standard space was performed through nonlinear registration. For functional data, we performed head motion correction before co-registering the functional data to the anatomical image with six degrees of freedom. Cerebrospinal fluid, white matter, and global signal, as well as head translation and rotation were extracted at each time point in the BOLD signal. We then used nilearn’s clean_img function to linearly detrend and apply a high-pass filter (0.01 Hz) to each functional time course [98]. We excluded any time courses with excessive motion parameters (motion > 3.0 mm or rotation > 3.0 degrees). Finally, we regressed out cerebrospinal fluid, white matter, and global signal, as well as the six head-motion parameters (three translations and three rotations), from the filtered functional data. To test how our results depend on global signal regression, we repeated all analyses without regressing out the global signal, leaving all other preprocessing steps unchanged.

### EEG preprocessing and spectral slope calculation

Brain Electrical Source Analysis software (BESA Research 7.1) was used to preprocess EEG recordings. We first removed fMRI artifacts from the EEG signal using BESA’s built-in functionality. Next, recordings were filtered using a 0.3 Hz high-pass filter, 80 Hz low-pass filter, and a 60 Hz notch filter to remove electrical noise. Voltages were then referenced to the average of all electrodes. Using tags corresponding to each fMRI TR, epochs were defined starting at 100 ms prior to TR and ending 2000 ms after TR. Voltage threshold detection was used for ocular artifact detection (voltage thresholds for the EOG electrodes were set at 150 μV for horizontal eye movements and 250 μV for vertical eye movements). Epochs were then additionally visually inspected by an expert for ocular artifacts including eye blinks and eye movements that were missed by the threshold detection. Identified artifacts were then removed from the epochs using ocular source components via BESA [99,100]. Certain artifacts, including sweat and myogenic artifacts (e.g., jaw clenching), could not be removed from the epochs using independent component analysis. Trials contaminated with these artifacts were not included in further analysis. Individual channels with excessive noise (greater than 150 μV), low signal (less than 0.01 μV), or excessive change (difference of 75 μV or more between adjacent channels) were replaced via interpolation using surrounding channels.

Spectral slopes were then calculated on the remaining epochs that did not have excessive noise. First, a power spectral density estimate was calculated using Welch’s method in SciPy (https://docs.scipy.org). Next, the power spectral density estimate was filtered from 0.5 Hz to 80 Hz, which was then log-transformed (using base 10). Finally, the slope of a fitted line to the log transformed spectral density plot was calculated as the point estimate for arousal during a specific epoch. This procedure was repeated for each epoch, yielding a time course of EEG spectral slopes.

### Pupillometry preprocessing

For the *Pupil Rest* pupil data, we used the preprocessed data provided by the authors [43], where the pupil time courses were blink-corrected, low-pass filtered, and downsampled to match the fMRI TR. For the *Pupil Story* pupil data, we followed the preprocessing procedures described in [46], where pupil time courses were similarly blink-corrected, interpolated, and downsampled. Each pupil time course was then z-scored within each run. For *Pupil Story*, the time courses were then averaged across participants to capture stimulus-evoked pupil fluctuations.

### Dynamic connectome-based predictive modeling

Whole-brain fMRI data were parcellated into 268 cortical and subcortical ROIs using the Shen atlas [61]. To assess whether our results depended on the choice of parcellation, we repeated all analyses using an alternative parcellation combining the 200-region Schaefer cortical parcellation [65] with 8 subcortical ROIs from the Brainnetome atlas [66] (bilateral amygdala, hippocampus, basal ganglia, and thalamus). Unlike the Shen atlas, the Schaefer-Brainnetome parcellation does not include the cerebellum.

For each ROI, we averaged the BOLD signal across voxels to obtain a single time course. For the resting-state and story-listening datasets, dynamic FC was computed using a tapered sliding window approach. Within each window, we calculated the Fisher z-transformed Pearson’s correlation between the time courses of every pair of ROIs (window size: *Pupil Rest*: 30 TRs = 30 s, *EEG Rest*: 14 TRs = 29.4 s, *Pupil Story*: 30 TRs = 30 s; Step size: 1 TR, Gaussian kernel σ = 7.5 s). The choice of window length involves a trade-off between temporal resolution and the reliability of functional connectivity estimates. Shorter windows can capture faster changes in arousal, but they estimate connectivity from fewer time points and are therefore noisier. We chose a window of approximately 30 s, similar to prior work using dynamic CPMs to predict fluctuations in psychological states [55,101]. To test whether our results depended on this choice, we repeated our analyses with 16-second and 44-second windows, again setting the standard deviation of the Gaussian kernel to one quarter of the window length. The EEG spectral slope and pupil dilation time courses were convolved with a hemodynamic response function and smoothed over the same Gaussian sliding window to match the fMRI data. For the movie datasets, we estimated FC within each narrative event to align with the event-level behavioral arousal ratings.

#### Within-dataset prediction

We trained dynamic CPMs using leave-one-participant-out cross-validation. In each fold, one participant served as the test set and the remaining *N -1* participants served as the training set. Feature selection was performed in the training set by correlating FC strength with the target arousal time course and retaining FCs that were positively or negatively correlated (two-tailed *t*-tests, *p* < 0.05). Significant positive and negative connections were assigned weights of +1 and -1, respectively, and all other connections were set to 0, yielding a binarized, signed arousal network for that fold. At each time point, we computed the element-wise product between the arousal network weights and the held-out participant’s dynamic FC matrix to obtain the predicted arousal time course.

Prediction accuracy was quantified as the Pearson correlation between the predicted and observed arousal time courses for the held-out participant. Correlation coefficients from all folds were Fisher *z*-transformed, averaged, and converted back to *r* values for reporting. Statistical significance was assessed using a non-parametric permutation test where the observed *r* was compared against a null distribution generated by repeating the analyses 10,000 times while phase-randomizing the target arousal time course and recomputing its correlation with the CPM-predicted time course, where *p* = (1 + # of null *r* ≥ observed *r*)/(1 + # of permutations).

#### Across-dataset prediction

To test generalization across datasets, we used the same dynamic CPM procedure described above, except that models were trained on all participants in one dataset to derive a single binarized, signed arousal network, and applied to each participant in an independent test dataset. When testing generalization from the movie datasets to the other datasets, we relied on a combined movies CPM trained on all 29 participants pooled across Sherlock and Film Festival, selecting edges with the same one-sample t-test across participants. Prediction accuracy was quantified as the Pearson correlation between predicted and target arousal time courses, with statistical significance assessed using the same phase-randomization permutation procedure described above. We calculated the generalization ratio for each prediction as the cross-dataset accuracy divided by the within-dataset accuracy for the test dataset.

#### Across-participant null

To account for temporal trends in arousal shared across participants, such as decreasing arousal with increasing scan duration [81,102,103], we constructed an additional null distribution for analyses in which Pupil Rest or EEG Rest was the test dataset. In each of 10,000 iterations, observed arousal time courses were randomly shuffled (i.e., reassigned) across participants. The correlation between each participant’s predicted time course and the reassigned observed time course was computed, Fisher z-transformed, and averaged across participants. Statistical significance was assessed by comparing the observed mean prediction accuracy against this null distribution. This analysis was not possible for the movie and story datasets, as their arousal time courses were derived from normative ratings and group-averaged pupil dilation and were therefore identical across participants.

### Network Overlap

We quantified the extent to which predictive connections were shared across the pupil-rest, EEG-rest, pupil-story, and combined movies CPMs by computing all two-way, three-way, and four-way overlap proportions. An edge was counted as overlapping only if it was selected as predictive in each CPM under consideration and showed the same direction of association with arousal (positive or negative). We assessed whether the observed overlap proportions exceeded chance using permutation testing. For each CPM, we randomly sampled the same number of positive and negative edges as in the corresponding empirical CPM, recomputed the overlap proportion for the relevant CPM combination, and repeated this procedure 1,000 times to generate a null distribution for each overlap statistic. *P*-values were computed as (1 + # of null proportions ≥ observed proportion)/(1 + # of permutations) [67].

We identified the subset of ROIs and edges comprising the four-way overlap set and summarized their distribution across large-scale functional networks. Since the Shen atlas ROIs do not map directly onto commonly used canonical functional network labels, we identified the Schaefer parcel with the greatest spatial overlap and assigned the Shen ROI the corresponding Yeo 7-network label associated with that Schaefer parcel [65]. Specifically, we used the 17-network version of the Schaefer parcellation and collapsed the 17 networks into seven by merging subnetworks within each network family and grouping the Temporal Parietal network with the Default Mode network [53]. This mapping facilitates comparison with prior work that uses Yeo network labels. ROIs in the cerebellum were assigned to a “Cerebellum” network, and subcortical ROIs were assigned to a “Subcortical” network.

To test whether the proportion of edges within or between specific network pairs exceeded what would be expected by chance, we compared the observed proportion against a null distribution. In each of 10,000 iterations, we randomly selected the same number of ROIs from the full Shen atlas as in the empirical four-way overlap set and randomly generated edges among those ROIs, matching the number of edges in the observed set. We computed the proportion of edges within and between network pairs in each iteration, and *p*-values were computed as (1 + # of null proportions ≥ observed proportion)/(1 + # of permutations). *P*-values were then corrected for multiple comparisons across all within-and between-network pairs using the Benjamini-Hochberg false discovery rate (FDR) procedure, and we report the resulting *q*-values.

We also constructed a CPM restricted to the edges shared across all four models (the four-way network). Each edge in the four-way overlap set was assigned a weight of +1 or -1 according to its shared direction of association with arousal, and all other edges were set to 0. With the Shen atlas, all shared edges were positive. We applied the four-way network to each dataset using the same procedure as for across-dataset prediction (**Figure S4A**). As the four-way overlap includes edges selected using the test dataset, we also tested a held-out version, in which each dataset was predicted from a network restricted to the edges shared by the other three models (**Figure S4B**).

### Similarity of Feature Maps

To compare the functional connectivity patterns associated with arousal in each dataset, we derived a feature map for each arousal CPM. The feature map at p < 0.05 was the binarized, signed arousal network obtained by training each CPM on all participants in the dataset, as in across-dataset prediction. To assess robustness, we also constructed feature maps thresholded at p < 0.01 and p < 0.10, as well as unthresholded maps in which each edge was represented by its mean Fisher z-transformed correlation with arousal across participants. For the sustained attention [51,68] and surprise [54] models, edges in each model’s positive network were assigned +1, edges in its negative network were assigned -1, and all other edges were assigned 0. We quantified the similarity between two models as the Pearson correlation between their feature maps across all edges.

As the sustained attention and surprise models were defined in the Shen atlas, we projected them into the Schaefer-Brainnetome parcellation to compare them with the arousal models in that parcellation. The projection was computed as WᵀMW, where M is the model’s signed 268 × 268 edge matrix and W gives the fraction of each Shen ROI’s voxels that falls within each Schaefer parcel. Edges involving the eight Brainnetome subcortical ROIs were set to 0.

To visualize the relationships between models, we embedded the pairwise similarities in two dimensions using interval multidimensional scaling (SMACOF; [104]) with distance defined as one minus the Pearson correlation. For each embedding, we retained the solution with the lowest stress-1 across a classical (Torgerson) start and 100 random starts. All embeddings were then aligned to the Shen *p* < 0.05 solution using Procrustes rotation to facilitate comparison.

### Bootstrap comparison of similarities

To test whether the arousal models were more similar to one another than to the comparison models, we computed the mean similarity among the four arousal models (six pairs), between the arousal models and the surprise model (four pairs), and between the arousal models and the sustained attention model (four pairs). We resampled participants with replacement within each dataset 10,000 times, rebuilt each arousal feature map from the resampled participants, and recomputed all similarities while holding the sustained attention and surprise models fixed. For each difference between the mean similarities, the one-sided *p*-value was computed as (1 + # of resamples in which the difference was ≤ 0)/(1 + # of resamples), and 95% confidence intervals were the 2.5th and 97.5th percentiles of the bootstrap distribution. The same participant draws were used for every feature-map type, parcellation, preprocessing choice, and window length.

### Predicting sleep stages from arousal CPMs

The Gu et al. (2023) simultaneous EEG-fMRI dataset [60] also includes data from several (1–8) 15-minute sleep sessions for each participant. The full sample included 33 participants, but we restricted our analyses to the same 24 participants used in the main analyses. Two additional participants had no usable sleep scans and were also excluded. EEG data were preprocessed to remove gradient and ballistocardiogram artifacts, re-referenced to the contralateral mastoid, and bandpass filtered at 0.3–35 Hz. A Registered Polysomnographic Technologist then scored each 30-second epoch as Wake, NREM1, NREM2, or NREM3 following AASM guidelines, as described in [105].

To generate a continuous target time course, we assigned ordinal values to each stage (Wake: 4, NREM1: 3, NREM2: 2, NREM3: 1), reflecting progressively deeper sleep and decreasing arousal. Dynamic FC was estimated within each 30-second epoch of the fMRI data to align with the temporal resolution of the sleep staging. We then applied each of the four arousal CPMs to predict the ordinal wakefulness time course. The pupil-rest, pupil-story, and movies CPMs were trained on participants independent of the EEG Sleep dataset and were applied to each participant’s sleep sessions using the same across-dataset procedure. We also applied a CPM containing the overlapping edges across all four models. As the EEG Rest and EEG Sleep data were acquired from the same participants, the EEG-rest CPM was evaluated using leave-one-participant-out cross-validation. For each participant, the EEG-rest CPM was trained on the resting-state data of the remaining participants and applied to that participant’s sleep sessions. Prediction accuracy and statistical significance were assessed using the same across-dataset procedure. Scans with only one sleep stage throughout were excluded from the analysis.

### Predicting memory from decoded arousal

Recall transcripts were obtained from the free recall task in the movie-watching datasets, in which participants verbally recounted the movie plot immediately after scanning. The segmentation and matching of recall transcripts to movie events were obtained from [57] for *Sherlock* and [46] for *Film Festival*. We embedded both event annotations and recall transcripts using Google’s Universal Sentence Encoder (USE) to obtain 512-dimensional vectors. USE embeds sentences with similar meaning in vectors closer in space [69]. Following prior work [46,53,70,106], we calculated “recall fidelity” as the cosine similarity between each event’s annotation vector and the participant’s recall vector for the corresponding event.

We tested whether CPM-decoded arousal during each event predicted later recall fidelity using Bayesian multilevel models implemented in *brms* [107]. For each CPM, decoded arousal was obtained at the event level by applying the corresponding arousal network to the movie functional connectivity data. This included CPMs trained on *Pupil Rest*, *EEG Rest*, and *Pupil Story*, as well as leave-one-out within-dataset predictions from the *Movies* datasets. We also applied the CPM containing the edges significant across all four datasets. Each model predicted recall fidelity from decoded arousal and included participant-level and dataset-level random intercepts, with both variables z-scored within each movie dataset.

We assumed weakly informative priors (see Supplemental Text). Models were fitted with four MCMC chains of 4,000 iterations each, discarding the first 1,000 iterations per chain as warm-up. Convergence was assessed using the Gelman–Rubin statistic (R^). We report posterior means and 95% highest density intervals (HDIs). Effects were considered credibly above zero when more than 95% of the posterior mass exceeded zero. Model fit was compared with a null model without the decoded arousal term using the Widely Applicable Information Criterion (WAIC), with lower values indicating better out-of-sample predictive performance.

### Brain Visualization

Brain visualizations were created using BrainNetViewer [108].

## Data Availability

All fMRI datasets used in this paper are publicly available on OpenNeuro. Arousal ratings for *Sherlock* and *Film Festival*, as well as pupil dilation data for *Pupil Story* are available at https://github.com/kannonb7/GeneralizedArousal. The predictive edges for each CPM, as well as the edges shared across CPMs, for both the Shen and Schaefer-Brainnetome parcellations are also available at the same repository.

## Code Availability

Analysis scripts are available at https://github.com/kannonb7/GeneralizedArousal.

## Acknowledgments

We thank all the authors who shared their data publicly and made this research possible. We acknowledge the University of Chicago’s Research Computing Center and the University of Chicago Social Sciences Division for their support of this work.

## Author Contributions

**K.B.:** Conceptualization, Data Curation, Methodology, Investigation, Software, Formal Analysis, Visualization, Writing -Original Draft, Writing -Review & Editing. **J.H.S.:** Conceptualization, Methodology, Investigation, Software, Formal Analysis, Writing -Original Draft. **Y.T.:** Software, Formal Analysis, Writing -Review & Editing. **Y.Z.:** Software, Formal Analysis. **J.K.:** Software, Writing -Review & Editing. **J.S.P.:** Software, Writing -Review & Editing. **A.C.:** Methodology, Software, Writing -Review & Editing. **M.D.R.:** Conceptualization, Methodology, Writing -Review & Editing, Supervision. **Y.C.L.:** Conceptualization, Writing -Original Draft, Writing -Review & Editing, Supervision, Project Administration.

