## Supplementary Information for "Situating affective, autonomic, and wakefulness arousal in a connectome-based neural reference space"

This PDF file includes:

Supplemental Text
Supplemental Figures S1–S7
Supplemental Tables S1–S6

**Supplemental Text**

**Priors for Bayesian multilevel models**

For each subject *i*, event *j*, and dataset *k*, the likelihood for an observed outcome $y_{i,j,k}$ is assumed to be selected from a normal distribution:

$$y_{i,j,k} \sim Normal(\mu_{i,j,k}, \sigma)$$

where $\mu_{i,j,k}$ is the expected outcome for subject *i*, event *j*, and dataset *k*, and $\sigma$ is the residual standard deviation of that outcome.

$\mu_{i,j,k}$ is modeled as a function of a fixed intercept and a fixed slope for the predictors with subject- and dataset-specific random intercepts.

$$\mu_{i,j,k} = \beta_0 + \alpha_{\text{subj[i]}} + \alpha_{\text{dataset[k]}} + \beta_1 x_{\text{i,j,k}}$$
$$\beta_0 \sim Normal(0, 1)$$
$$\beta_1 \sim Normal(0, 1)$$
$$\alpha_{\text{subj[i]}} \sim Normal(0, \sigma_{\text{subj}})$$
$$\alpha_{\text{dataset[k]}} \sim Normal(0, \sigma_{\text{dataset}})$$
$$\sigma_{\text{subj}} \sim Exponential(1)$$
$$\sigma_{\text{dataset}} \sim Exponential(1)$$
$$\sigma \sim Exponential(1)$$

We used weakly informative priors. Fixed effects were assigned *Normal*(0, 1) priors centered at zero, with no strong assumptions about effect direction while allowing reasonable effect sizes. The standard deviation of 1 provides mild regularization to reduce overfitting without imposing strong constraints. Standard deviation parameters were given *Exponential*(1) priors, which restrict values to be positive and favor smaller variance.

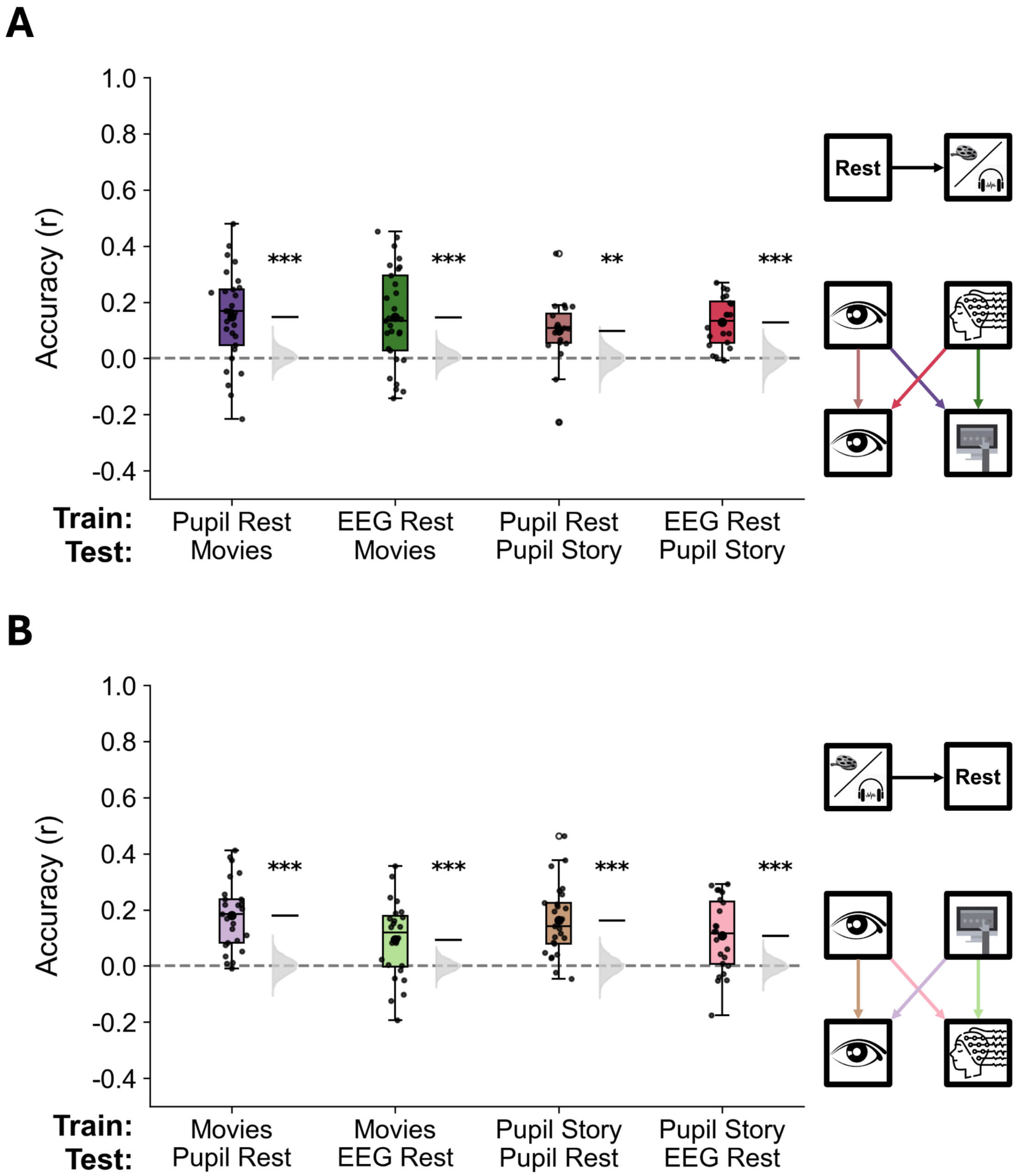


**Figure S1. Across-dataset prediction accuracy when dynamic functional connectivity is calculated over a 16-second sliding window.** CPM performance in predicting arousal across datasets for pupil dilation (*Pupil Rest*), EEG spectral slope (*EEG Rest*), behavioral arousal ratings during movie-watching (*Movies*), and pupil dilation during story listening (*Pupil Story*). The y-axis displays the prediction accuracy, as measured by Pearson's correlation between the model's predicted time course and the observed arousal time course. Each datapoint within the box plot denotes a participant. The black horizontal lines show the average of the Fisher z-transformed r-values. The gray half-violin plots show the null distributions generated from correlating the predicted arousal time courses with phase-randomized observed arousal time courses. *$p < .05$, ** $p < .01$, *** $p < .001$.

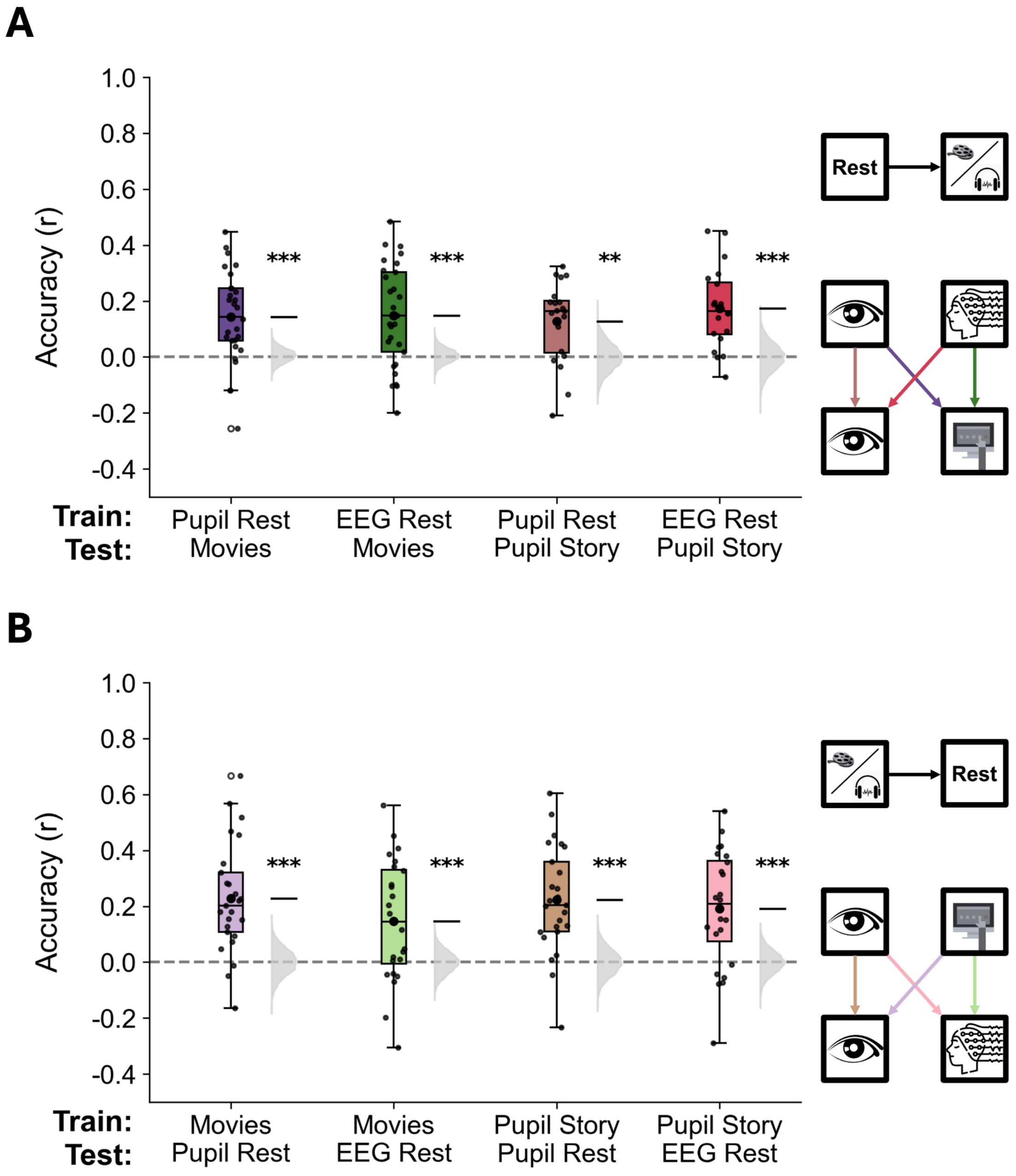


**Figure S2. Across-dataset prediction accuracy when dynamic functional connectivity is calculated over a 44-second sliding window.** CPM performance in predicting arousal across datasets for pupil dilation (*Pupil Rest*), EEG spectral slope (*EEG Rest*), behavioral arousal ratings during movie-watching (*Movies*), and pupil dilation during story listening (*Pupil Story*). The y-axis displays the prediction accuracy, as measured by Pearson's correlation between the model's predicted time course and the observed arousal time course. Each datapoint within the box plot denotes a participant. The black horizontal lines show the average of the Fisher z-transformed r-values. The gray half-violin plots show the null distributions generated from correlating the predicted arousal time courses with phase-randomized observed arousal time courses. *$p < .05$, ** $p < .01$, *** $p < .001$.

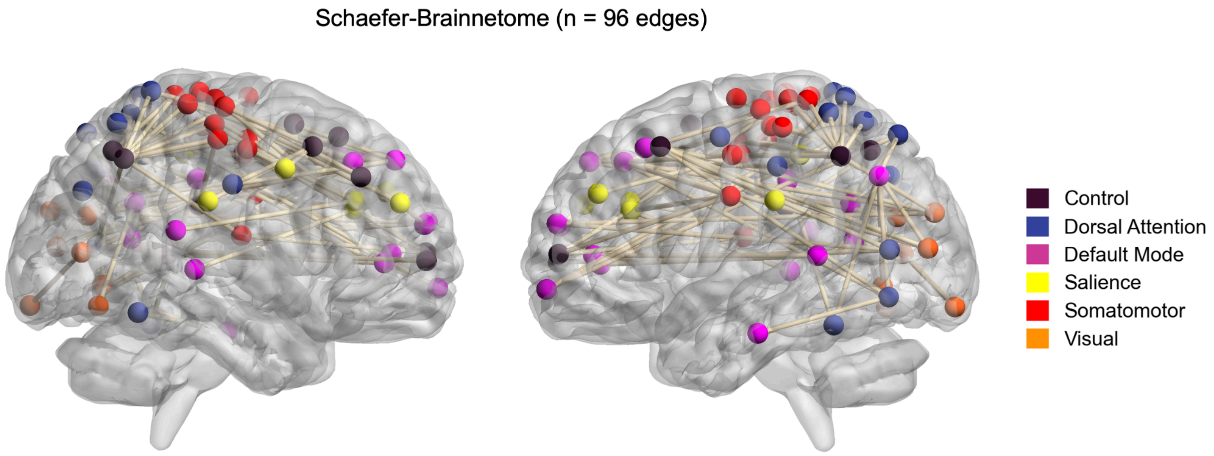


**Figure S3. Glass brain displaying 96 functional connections that overlap across all four CPMs using the Schaefer-Brainnetome atlas.** Nodes indicate brain regions, with colors denoting functional networks. Limbic and subcortical nodes were not present in the four-way overlap network. The Schaefer-Brainnetome atlas does not include the cerebellum.

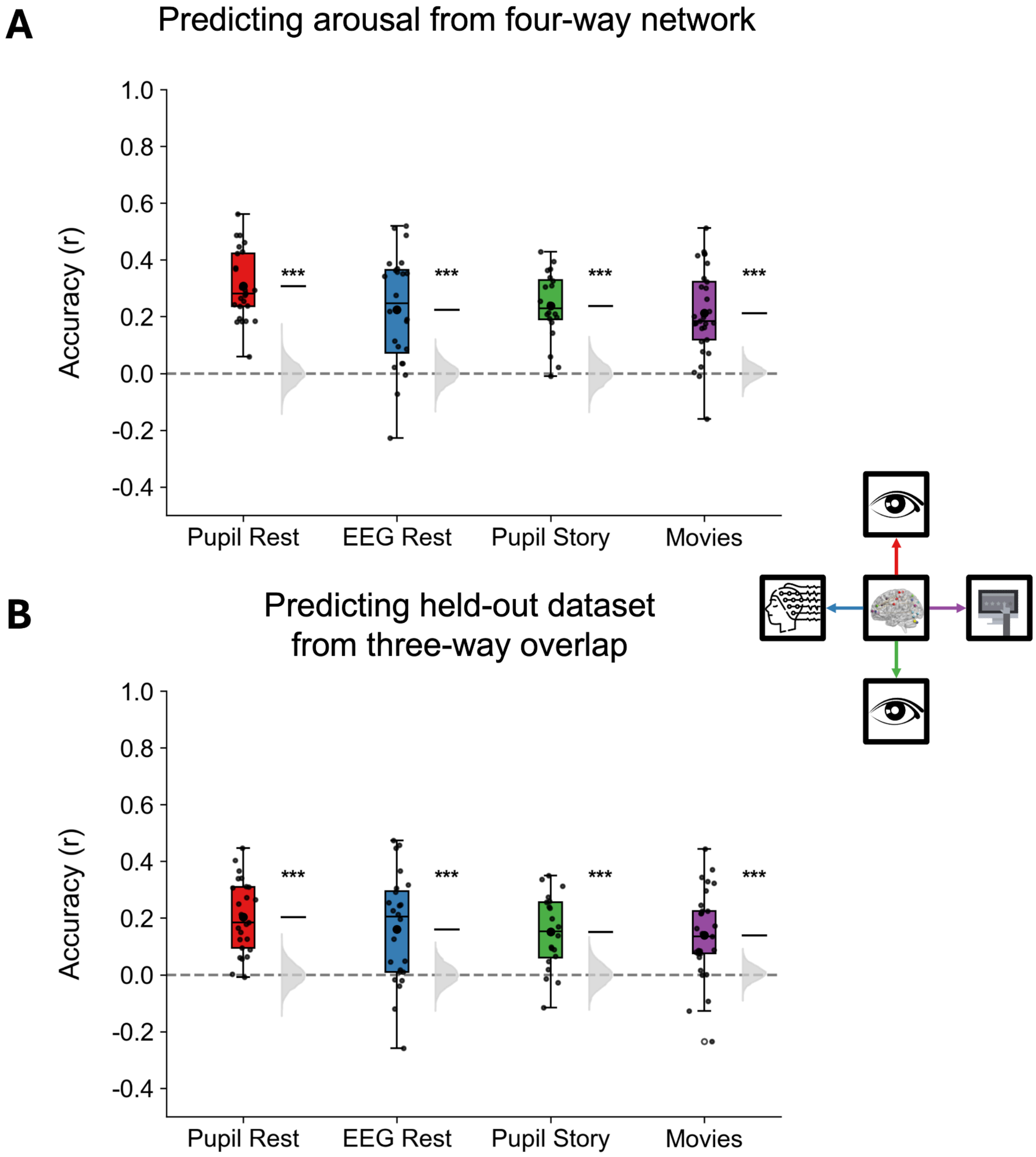


**Figure S4. Dynamic CPM trained on overlapping arousal networks generalizes to all datasets.** CPM performance in predicting arousal across datasets for pupil dilation (*Pupil Rest*), EEG spectral slope (*EEG Rest*), behavioral arousal ratings during movie-watching (*Movies*), and pupil dilation during story listening (*Pupil Story*). **(A)** A CPM restricted to features shared across all four models significantly predicted arousal fluctuations in all datasets. **(B)** A CPM restricted to edges shared by the other three models significantly predicted arousal fluctuations in the held-out dataset. The y-axis displays model prediction accuracy, as measured by Pearson's correlation between the model's predicted time course and the observed arousal time course. Each datapoint within the box plot denotes a participant. The black horizontal lines show the average of the Fisher z-transformed *r*-values. The gray half-violin plots show the null distributions generated from correlating the predicted arousal time courses with phase-randomized observed arousal time courses. *$p < .05$, ** $p < .01$, *** $p < .001$.

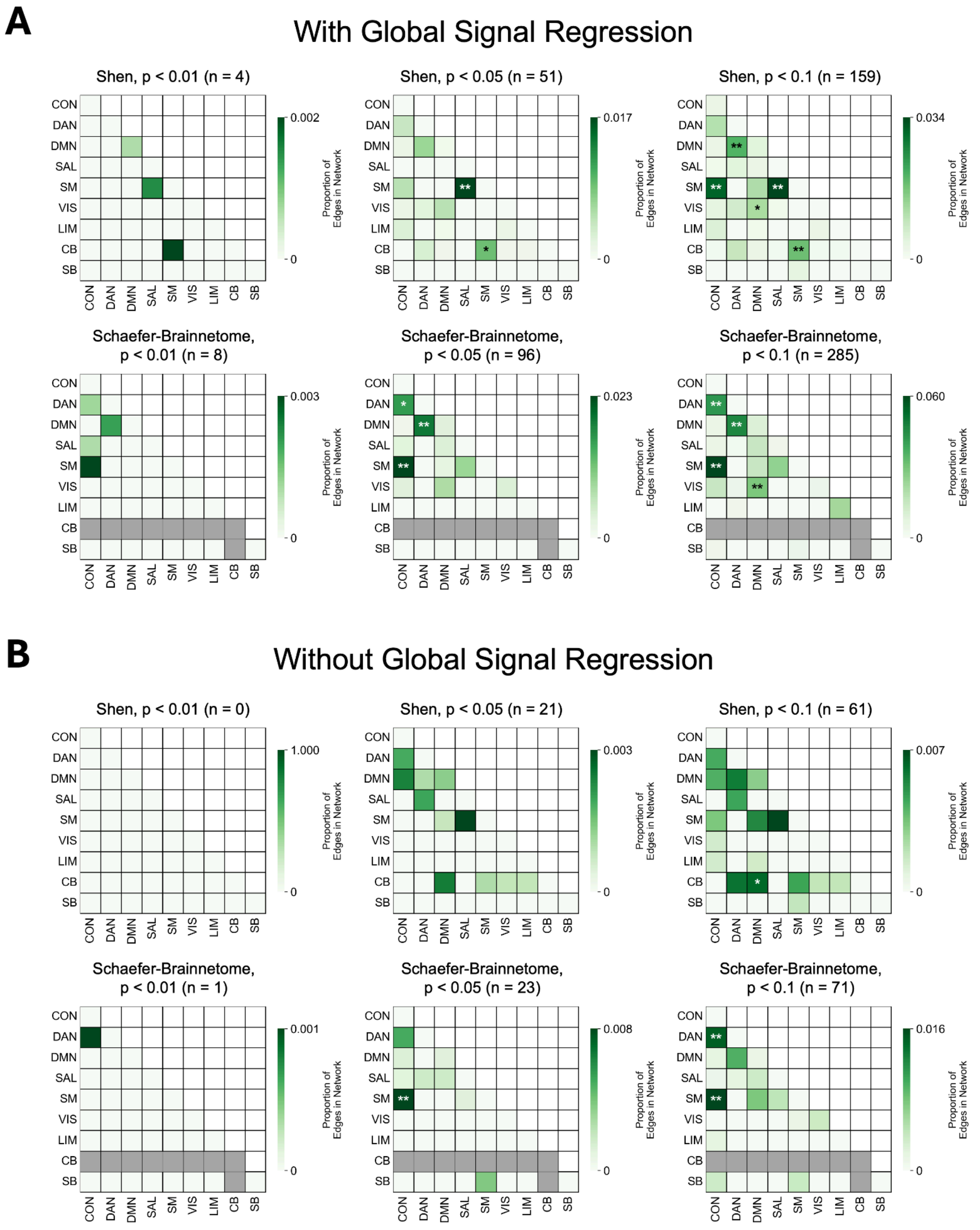


**Figure S5. Proportion of edges within and between each pair of functional networks common across all four models. (A)** Global signal regression applied during preprocessing. **(B)** Global signal regression not applied during preprocessing. Columns show feature-selection thresholds, and rows show the Shen and Schaefer-Brainnetome parcellations, with the number of shared edges given in each panel title. The Schaefer-Brainnetome atlas does not include the cerebellum. CON: Control, DAN: Dorsal Attention, DMN: Default Mode, SAL: Salience, SM: Somatomotor, VIS: Visual, LIM: Limbic, CB: Cerebellum, SB: Subcortical. * q < 0.05, ** q < 0.01, *** q < 0.001.

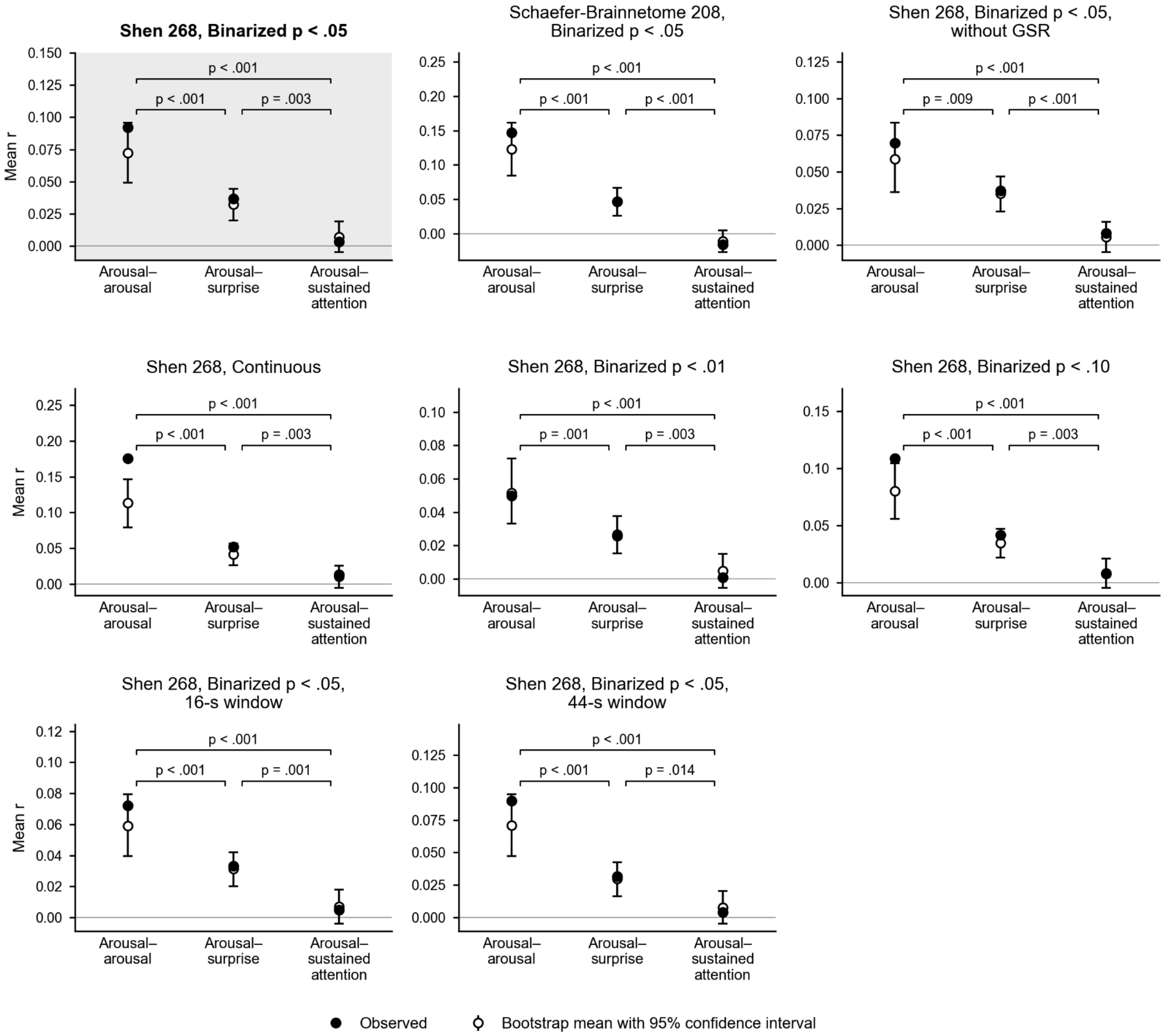


**Figure S6. Comparing similarity of feature maps across models.** Each panel shows the mean pairwise correlation between feature maps for pairs of arousal models (arousal-arousal), and for each arousal model paired with the surprise model (arousal-surprise) or the sustained attention model (arousal-sustained attention). Filled circles show observed values, and open circles with error bars show bootstrap means and 95% confidence intervals from resampling participants. Bootstrap means are lower than observed values because resampling participants adds noise to each arousal model's feature map, which attenuates correlations between maps. This attenuation is strongest for arousal-arousal pairs, so the bootstrap test provides a conservative test of differences between the arousal models and the comparison models. Brackets show bootstrap p-values for differences in mean similarity. The shaded panel shows the main analysis reported in the main text. The other panels show the same analysis with different feature-selection criteria ($p < 0.01$, $p < 0.05$, $p < 0.10$, continuous), sliding-window length (16 s, 44 s), a different parcellation (Shen, Schaefer-Brainnetome), or without global signal regression.

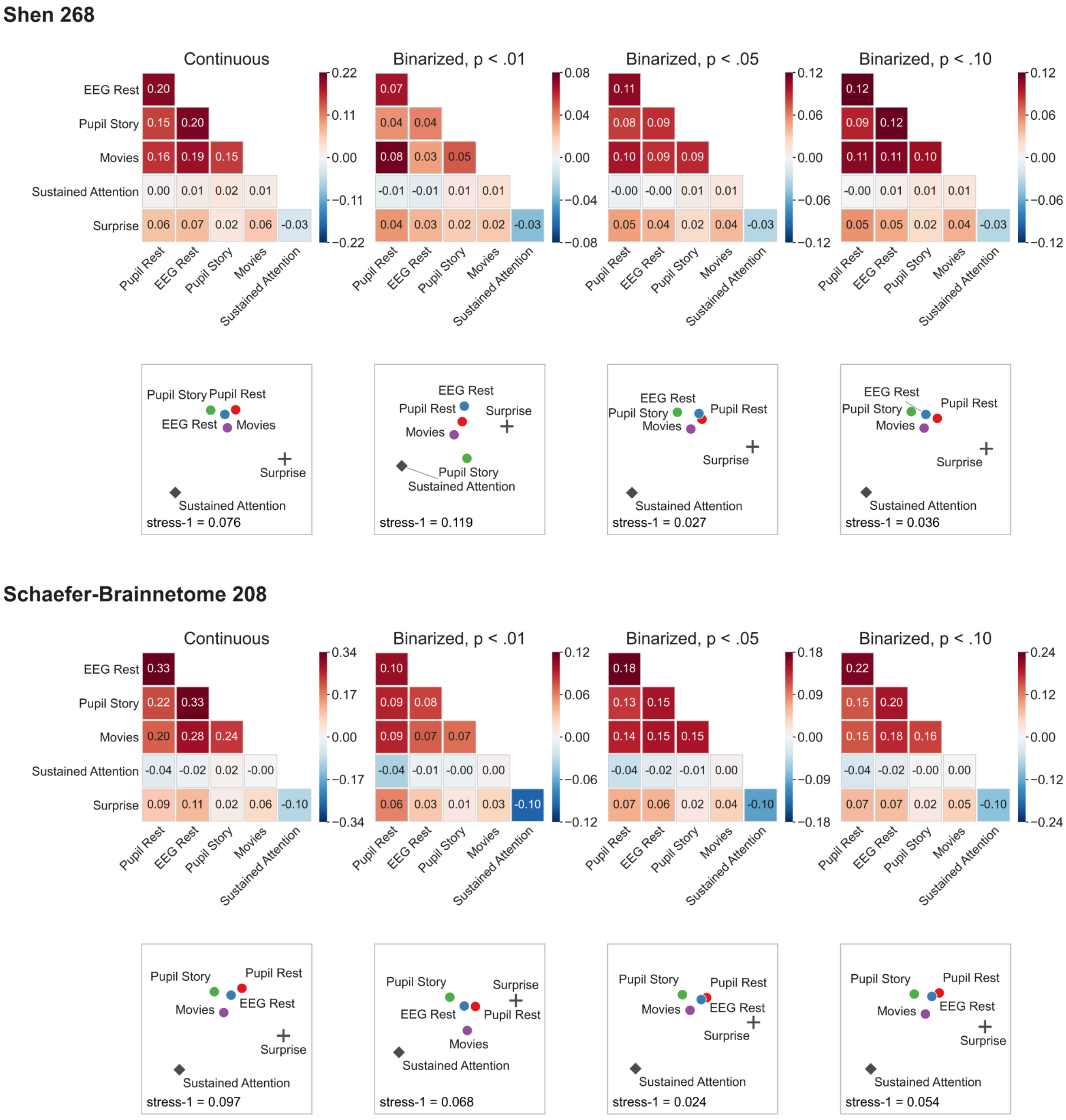


**Figure S7. Pairwise similarity of feature maps across models.** Results are shown for the Shen 268 parcellation (top) and the Schaefer-Brainnetome 208 parcellation (bottom), using continuous feature maps and feature maps binarized at $p < 0.01$, $p < 0.05$, and $p < 0.10$. For each condition, the matrix shows the Pearson correlation between the feature maps of each pair of models, and the panel below it shows a two-dimensional embedding of these similarities obtained with multidimensional scaling, with distance defined as one minus the correlation. Stress-1 values indicate how well each embedding preserves the pairwise distances, with lower values indicating a better fit. For the Schaefer-Brainnetome parcellation, the comparison models were projected from the Shen atlas. All analyses used global signal regression and a 30-second sliding window. The Shen 268, $p < 0.05$ embedding corresponds to Figure 6C.

**Table S1. Prediction accuracy across parcellation schemes.** Model prediction accuracy for CPMs constructed with the Shen 268-region atlas and with the Schaefer 200-region cortical parcellation combined with 8 Brainnetome subcortical regions. Generalization ratios of cross-dataset predictions are computed by dividing the cross-dataset accuracy by the within-dataset accuracy.

| Train | Test | Shen *r* | Shen *p* | Generalization Ratio | Schaefer + Brainnetome r | Schaefer + Brainnetome p | Generalization Ratio |
|---|---|---|---|---|---|---|---|
| *Within-dataset* | | | | | | | |
| Pupil Rest | Pupil Rest | 0.308 | <0.001 | - | 0.278 | <0.001 | - |
| EEG Rest | EEG Rest | 0.201 | <0.001 | - | 0.202 | <0.001 | - |
| Pupil Story | Pupil Story | 0.250 | <0.001 | - | 0.248 | <0.001 | - |
| Movies | Movies | 0.340 | <0.001 | - | 0.369 | <0.001 | - |
| *Across rest datasets* | | | | | | | |
| Pupil Rest | EEG Rest | 0.150 | <0.001 | 0.746 | 0.165 | <0.001 | 0.817 |
| EEG Rest | Pupil Rest | 0.243 | <0.001 | 0.789 | 0.240 | <0.001 | 0.863 |
| *Across narrative datasets* | | | | | | | |
| Movies | Pupil Story | 0.136 | <0.001 | 0.544 | 0.159 | <0.001 | 0.641 |
| Pupil Story | Movies | 0.162 | <0.001 | 0.476 | 0.163 | <0.001 | 0.442 |
| *Rest to narrative* | | | | | | | |
| Pupil Rest | Movies | 0.151 | <0.001 | 0.444 | 0.139 | <0.001 | 0.377 |
| EEG Rest | Movies | 0.161 | <0.001 | 0.474 | 0.164 | <0.001 | 0.444 |
| Pupil Rest | Pupil Story | 0.121 | 0.001 | 0.484 | 0.144 | <0.001 | 0.581 |
| EEG Rest | Pupil Story | 0.169 | <0.001 | 0.676 | 0.179 | <0.001 | 0.722 |
| *Narrative to rest* | | | | | | | |
| Movies | Pupil Rest | 0.230 | <0.001 | 0.747 | 0.199 | <0.001 | 0.716 |
| Movies | EEG Rest | 0.148 | <0.001 | 0.736 | 0.158 | <0.001 | 0.782 |
| Pupil Story | Pupil Rest | 0.215 | <0.001 | 0.698 | 0.185 | <0.001 | 0.665 |
| Pupil Story | EEG Rest | 0.179 | <0.001 | 0.891 | 0.166 | <0.001 | 0.822 |

**Table S2. Prediction accuracy across sliding windows.** Model prediction accuracy for CPMs using the Shen atlas parcellation when computing dynamic functional connectivity with sliding windows of 16 or 44 seconds.

| Train | Test | 16-second *r* | 16-second *p* | 44-second *r* | 44-second *p* |
|---|---|---|---|---|---|
| *Within-dataset* | | | | | |
| Pupil Rest | Pupil Rest | 0.280 | <0.001 | 0.308 | <0.001 |
| EEG Rest | EEG Rest | 0.149 | <0.001 | 0.214 | <0.001 |
| Pupil Story | Pupil Story | 0.198 | <0.001 | 0.274 | <0.001 |
| Movies | Movies | 0.340[1] | <0.001 | 0.340[1] | <0.001 |
| *Across rest datasets* | | | | | |
| Pupil Rest | EEG Rest | 0.105 | <0.001 | 0.157 | <0.001 |
| EEG Rest | Pupil Rest | 0.186 | <0.001 | 0.255 | <0.001 |
| *Across narrative datasets* | | | | | |
| Movies | Pupil Story | 0.098 | <0.001 | 0.159 | <0.001 |
| Pupil Story | Movies | 0.134 | <0.001 | 0.183 | <0.001 |
| *Rest to narrative* | | | | | |
| Pupil Rest | Movies | 0.152 | <0.001 | 0.147 | <0.001 |
| EEG Rest | Movies | 0.150 | <0.001 | 0.154 | <0.001 |
| Pupil Rest | Pupil Story | 0.099 | 0.001 | 0.129 | 0.003 |
| EEG Rest | Pupil Story | 0.130 | <0.001 | 0.178 | <0.001 |
| *Narrative to rest* | | | | | |
| Movies | Pupil Rest | 0.183 | <0.001 | 0.239 | <0.001 |
| Movies | EEG Rest | 0.095 | <0.001 | 0.154 | <0.001 |
| Pupil Story | Pupil Rest | 0.165 | <0.001 | 0.232 | <0.001 |
| Pupil Story | EEG Rest | 0.109 | <0.001 | 0.199 | <0.001 |

[1]Within-dataset Movies predictions were computed at the event level, so these numbers are consistent across sliding windows.

**Table S3. Prediction accuracy when the global signal is not regressed from fMRI data.** Model prediction accuracy for CPMs constructed without global signal regression with the Shen 268-region atlas and with the Schaefer 200-region cortical parcellation combined with 8 Brainnetome subcortical regions.

| Train | Test | Shen *r* | Shen *p* | Schaefer + Brainnetome r | Schaefer + Brainnetome p |
|---|---|---|---|---|---|
| *Within-dataset* | | | | | |
| Pupil Rest | Pupil Rest | 0.206 | <0.001 | 0.192 | <0.001 |
| EEG Rest | EEG Rest | 0.161 | <0.001 | 0.132 | <0.001 |
| Pupil Story | Pupil Story | 0.207 | <0.001 | 0.226 | <0.001 |
| Movies | Movies | 0.301 | <0.001 | 0.333 | <0.001 |
| *Across rest datasets* | | | | | |
| Pupil Rest | EEG Rest | 0.092 | 0.002 | 0.087 | 0.003 |
| EEG Rest | Pupil Rest | 0.218 | <0.001 | 0.222 | <0.001 |
| *Across narrative datasets* | | | | | |
| Movies | Pupil Story | 0.072 | 0.028 | 0.112 | <0.001 |
| Pupil Story | Movies | 0.069 | 0.005 | 0.112 | <0.001 |
| *Rest to narrative* | | | | | |
| Pupil Rest | Movies | 0.058 | 0.017 | 0.055 | 0.026 |
| EEG Rest | Movies | 0.045 | 0.049 | 0.077 | 0.002 |
| Pupil Rest | Pupil Story | 0.008 | 0.409 | 0.020 | 0.297 |
| EEG Rest | Pupil Story | 0.029 | 0.207 | 0.075 | 0.016 |
| *Narrative to rest* | | | | | |
| Movies | Pupil Rest | 0.222 | <0.001 | 0.159 | <0.001 |
| Movies | EEG Rest | 0.094 | 0.001 | 0.108 | <0.001 |
| Pupil Story | Pupil Rest | 0.025 | 0.276 | 0.067 | 0.039 |
| Pupil Story | EEG Rest | 0.058 | 0.033 | 0.129 | <0.001 |

**Table S4. Table of common set of predictive edges across all datasets when using the Shen atlas.** All edges in this table are positive.

| Network | Shen ROI # | Network | Shen ROI # |
|---|---|---|---|
| Default | 5 | Dorsal Attention | 209 |
| Salience | 9 | Somatomotor | 158 |
| Default | 10 | Visual | 208 |
| Default | 10 | Control | 221 |
| Default | 12 | Dorsal Attention | 32 |
| Default | 12 | Cerebellum | 236 |
| Control | 14 | Somatomotor | 33 |
| Control | 14 | Somatomotor | 39 |
| Control | 14 | Dorsal Attention | 45 |
| Control | 14 | Limbic | 71 |
| Somatomotor | 23 | Control | 142 |
| Somatomotor | 24 | Cerebellum | 116 |
| Somatomotor | 25 | Salience | 219 |
| Somatomotor | 27 | Salience | 219 |
| Somatomotor | 33 | Cerebellum | 102 |
| Somatomotor | 33 | Salience | 150 |
| Somatomotor | 33 | Salience | 219 |
| Somatomotor | 39 | Cerebellum | 100 |
| Somatomotor | 39 | Cerebellum | 102 |
| Somatomotor | 39 | Salience | 144 |
| Somatomotor | 39 | Salience | 146 |
| Control | 48 | Limbic | 71 |
| Control | 55 | Visual | 81 |
| Limbic | 56 | Visual | 81 |
| Dorsal Attention | 66 | Visual | 73 |
| Visual | 67 | Default | 148 |
| Limbic | 71 | Cerebellum | 107 |
| Limbic | 71 | Default | 182 |
| Visual | 81 | Cerebellum | 100 |

| | | | |
|---|---|---|---|
| Cerebellum | 102 | Somatomotor | 174 |
| Cerebellum | 114 | Dorsal Attention | 206 |
| Cerebellum | 116 | Somatomotor | 174 |
| Control | 143 | Dorsal Attention | 171 |
| Salience | 144 | Somatomotor | 158 |
| Default | 148 | Visual | 208 |
| Default | 148 | Dorsal Attention | 209 |
| Default | 149 | Dorsal Attention | 206 |
| Default | 149 | Visual | 208 |
| Default | 149 | Visual | 210 |
| Somatomotor | 158 | Salience | 219 |
| Somatomotor | 159 | Salience | 219 |
| Somatomotor | 159 | Cerebellum | 247 |
| Somatomotor | 174 | Cerebellum | 241 |
| Dorsal Attention | 177 | Default | 182 |
| Control | 178 | Default | 225 |
| Default | 182 | Default | 203 |
| Default | 182 | Dorsal Attention | 206 |
| Default | 182 | Visual | 208 |
| Default | 182 | Visual | 210 |
| Default | 192 | Cerebellum | 246 |
| Dorsal Attention | 206 | Cerebellum | 247 |

**Table S5. Table of common set of predictive edges across all datasets when using the combined Schaefer-Brainnetome Atlas.** Pos and neg indicate positive and negative edges respectively.

| ROI | Network | ROI # | ROI | Network | ROI # | Sign |
|---|---|---|---|---|---|---|
| L extrastriate cortex | Visual | 2 | R striate/calcarine cortex | Visual | 109 | neg |
| L extrastriate cortex | Visual | 5 | L lateral prefrontal cortex | Control | 69 | pos |
| L extrastriate cortex | Visual | 5 | L inferior parietal lobule | Default | 75 | pos |
| L extrastriate cortex | Visual | 5 | L dorsal prefrontal cortex | Default | 76 | pos |
| L extrastriate cortex | Visual | 5 | L dorsal prefrontal cortex | Default | 90 | pos |
| L extrastriate cortex | Visual | 6 | L inferior parietal lobule | Control | 68 | pos |
| L extrastriate cortex | Visual | 6 | L lateral prefrontal cortex | Control | 69 | pos |
| L extrastriate cortex | Visual | 6 | L dorsal prefrontal cortex | Default | 88 | pos |
| L extrastriate cortex | Visual | 6 | L dorsal prefrontal cortex | Default | 90 | pos |
| L inferior extrastriate cortex | Visual | 8 | L inferior parietal lobule | Default | 75 | pos |
| L superior extrastriate cortex | Visual | 11 | R dorsal prefrontal cortex | Default | 192 | pos |
| L somatomotor cortex | Somatomotor | 13 | L lateral prefrontal cortex | Salience | 48 | pos |
| L somatomotor cortex | Somatomotor | 13 | L inferior parietal lobule | Control | 68 | pos |
| L somatomotor cortex | Somatomotor | 14 | R intraparietal sulcus | Control | 165 | pos |
| L somatomotor cortex | Somatomotor | 15 | R intraparietal sulcus | Control | 165 | pos |
| L somatomotor cortex | Somatomotor | 16 | R dorsolateral prefrontal cortex | Control | 175 | pos |
| L somatomotor cortex | Somatomotor | 17 | L inferior parietal lobule | Control | 68 | pos |
| L somatomotor cortex | Somatomotor | 17 | L dorsal prefrontal cortex | Default | 76 | pos |
| L somatomotor cortex | Somatomotor | 18 | R inferior parietal lobule | Control | 173 | pos |
| L somatomotor cortex | Somatomotor | 19 | L inferior parietal lobule | Control | 68 | pos |
| L somatomotor cortex | Somatomotor | 19 | R lateral prefrontal cortex | Salience | 153 | pos |
| L somatomotor cortex | Somatomotor | 19 | R posterior medial prefrontal cortex | Salience | 156 | pos |
| L somatomotor cortex | Somatomotor | 19 | R inferior parietal lobule | Control | 173 | pos |
| L somatomotor cortex | Somatomotor | 19 | R dorsolateral prefrontal cortex | Control | 175 | pos |
| L somatomotor cortex | Somatomotor | 20 | L inferior parietal lobule | Control | 68 | pos |
| L central sulcus | Somatomotor | 27 | L posterior medial prefrontal cortex | Salience | 50 | pos |
| L central sulcus | Somatomotor | 27 | L ventrolateral prefrontal cortex | Control | 71 | pos |
| L central sulcus | Somatomotor | 27 | L dorsal prefrontal cortex | Default | 90 | pos |
| L central sulcus | Somatomotor | 28 | L dorsal prefrontal cortex | Default | 90 | pos |
| L temporo-occipital cortex | Dorsal Attention | 29 | L inferior parietal lobule | Default | 75 | pos |

| | | | | | | |
|---|---|---|---|---|---|---|
| L temporo-occipital cortex | Dorsal Attention | 29 | L precuneus/posterior cingulate cortex | Default | 78 | pos |
| L temporo-occipital cortex | Dorsal Attention | 30 | L lateral prefrontal cortex | Control | 69 | pos |
| L temporo-occipital cortex | Dorsal Attention | 30 | L inferior parietal lobule | Default | 75 | pos |
| L temporo-occipital cortex | Dorsal Attention | 30 | L temporal cortex | Default | 84 | pos |
| L temporo-occipital cortex | Dorsal Attention | 30 | L dorsal prefrontal cortex | Default | 90 | pos |
| L parieto-occipital cortex | Dorsal Attention | 31 | L lateral prefrontal cortex | Control | 69 | pos |
| L parieto-occipital cortex | Dorsal Attention | 31 | L dorsal prefrontal cortex | Default | 88 | pos |
| L parieto-occipital cortex | Dorsal Attention | 31 | R inferior parietal lobule | Control | 173 | pos |
| L parieto-occipital cortex | Dorsal Attention | 31 | R ventrolateral prefrontal cortex | Control | 178 | pos |
| L parieto-occipital cortex | Dorsal Attention | 31 | R medial prefrontal cortex | Default | 187 | pos |
| L superior parietal lobule | Dorsal Attention | 32 | L inferior parietal lobule | Default | 75 | pos |
| L superior parietal lobule | Dorsal Attention | 33 | L inferior parietal lobule | Default | 75 | pos |
| L superior parietal lobule | Dorsal Attention | 33 | L precuneus/posterior cingulate cortex | Default | 77 | pos |
| L superior parietal lobule | Dorsal Attention | 33 | L medial prefrontal cortex | Default | 81 | pos |
| L superior parietal lobule | Dorsal Attention | 34 | L inferior parietal lobule | Control | 68 | pos |
| L postcentral gyrus | Dorsal Attention | 35 | R dorsal prefrontal cortex | Default | 185 | pos |
| L postcentral gyrus | Dorsal Attention | 37 | L inferior parietal lobule | Control | 68 | pos |
| L postcentral gyrus | Dorsal Attention | 38 | L inferior parietal lobule | Control | 68 | pos |
| L postcentral gyrus | Dorsal Attention | 38 | R inferior parietal lobule | Control | 173 | pos |
| L frontal eye field | Dorsal Attention | 39 | R dorsal prefrontal cortex | Default | 185 | pos |
| L parietal operculum | Salience | 40 | R dorsolateral prefrontal cortex | Control | 180 | pos |
| L medial parietal cortex | Salience | 44 | R dorsal prefrontal cortex | Default | 192 | pos |
| L posterior medial prefrontal cortex | Salience | 50 | R somatomotor cortex | Somatomotor | 114 | pos |
| L posterior medial prefrontal cortex | Salience | 50 | R somatomotor cortex | Somatomotor | 118 | pos |
| L posterior medial prefrontal cortex | Salience | 50 | R temporoparietal junction | Default | 198 | pos |
| L intraparietal sulcus | Control | 58 | R somatomotor cortex | Somatomotor | 121 | pos |
| L inferior parietal lobule | Control | 68 | R somatomotor cortex | Somatomotor | 115 | pos |
| L inferior parietal lobule | Control | 68 | R somatomotor cortex | Somatomotor | 119 | pos |
| L inferior parietal lobule | Control | 68 | R somatomotor cortex | Somatomotor | 121 | pos |
| L inferior parietal lobule | Control | 68 | R somatomotor cortex | Somatomotor | 123 | pos |
| L inferior parietal lobule | Control | 68 | R superior parietal lobule | Dorsal Attention | 136 | pos |
| L inferior parietal lobule | Control | 68 | R postcentral gyrus | Dorsal Attention | 140 | pos |

| | | | | | | |
|---|---|---|---|---|---|---|
| L ventrolateral prefrontal cortex | Control | 71 | R somatomotor cortex | Somatomotor | 116 | pos |
| L ventrolateral prefrontal cortex | Control | 71 | R secondary somatosensory cortex | Somatomotor | 126 | pos |
| L precuneus | Control | 73 | L medial prefrontal cortex | Default | 81 | pos |
| L posterior cingulate cortex | Control | 74 | R somatomotor cortex | Somatomotor | 118 | pos |
| L inferior parietal lobule | Default | 75 | L retrosplenial cortex | Default | 97 | pos |
| L inferior parietal lobule | Default | 75 | R superior parietal lobule | Dorsal Attention | 133 | pos |
| L precuneus/posterior cingulate cortex | Default | 77 | R extrastriate cortex | Visual | 101 | pos |
| L medial prefrontal cortex | Default | 81 | R postcentral gyrus | Dorsal Attention | 139 | pos |
| L medial prefrontal cortex | Default | 82 | L temporoparietal junction | Default | 99 | pos |
| L temporal cortex | Default | 84 | R temporo-occipital cortex | Dorsal Attention | 131 | pos |
| L dorsal prefrontal cortex | Default | 90 | R somatomotor cortex | Somatomotor | 119 | pos |
| L temporoparietal junction | Default | 99 | R medial prefrontal cortex | Default | 187 | pos |
| R somatomotor cortex | Somatomotor | 113 | R inferior parietal lobule | Control | 173 | pos |
| R somatomotor cortex | Somatomotor | 114 | R posterior medial prefrontal cortex | Salience | 156 | pos |
| R somatomotor cortex | Somatomotor | 115 | R intraparietal sulcus | Control | 165 | pos |
| R somatomotor cortex | Somatomotor | 115 | R inferior parietal lobule | Control | 173 | pos |
| R somatomotor cortex | Somatomotor | 116 | R inferior parietal lobule | Control | 173 | pos |
| R somatomotor cortex | Somatomotor | 118 | R dorsolateral prefrontal cortex | Control | 176 | pos |
| R somatomotor cortex | Somatomotor | 121 | R posterior medial prefrontal cortex | Salience | 156 | pos |
| R somatomotor cortex | Somatomotor | 121 | R intraparietal sulcus | Control | 165 | pos |
| R somatomotor cortex | Somatomotor | 121 | R inferior parietal lobule | Control | 173 | pos |
| R somatomotor cortex | Somatomotor | 122 | R dorsolateral prefrontal cortex | Control | 175 | pos |
| R somatomotor cortex | Somatomotor | 123 | R inferior parietal lobule | Control | 173 | pos |
| R secondary somatosensory cortex | Somatomotor | 126 | R inferior parietal lobule | Control | 173 | pos |
| R postcentral gyrus | Dorsal Attention | 137 | R dorsolateral prefrontal cortex | Control | 176 | pos |
| R postcentral gyrus | Dorsal Attention | 137 | R dorsal prefrontal cortex | Default | 192 | pos |
| R postcentral gyrus | Dorsal Attention | 139 | R inferior parietal lobule | Control | 173 | pos |
| R postcentral gyrus | Dorsal Attention | 140 | R inferior parietal lobule | Control | 173 | pos |
| R postcentral gyrus | Dorsal Attention | 140 | R dorsal prefrontal cortex | Default | 192 | pos |
| R parietal operculum | Salience | 142 | R dorsolateral prefrontal cortex | Control | 176 | pos |
| R precentral cortex | Salience | 143 | R dorsolateral prefrontal | Control | 176 | pos |

| | | | | | | |
|---|---|---|---|---|---|---|
| | | | cortex | | | |
| R lateral prefrontal cortex | Salience | 153 | R medial prefrontal cortex | Default | 189 | pos |
| R posterior medial prefrontal cortex | Salience | 156 | R temporoparietal junction | Default | 200 | pos |
| R dorsal prefrontal cortex | Control | 167 | R dorsal prefrontal cortex | Default | 192 | pos |

**Table S6. Subcortical edges were shared across combinations of three arousal networks.** Using the 8 Brainnetome subcortical ROIs that we appended to the Schaefer-200 Atlas, we examined the subcortical ROIs represented in the common edges across every combination of three out of the four datasets. Edges containing subcortical ROIs were present in each three-way network.

| Three-Way Network | Total Number of Edges | Number of Subcortical Edges | Subcortical ROIs |
|---|---|---|---|
| *Pupil Rest*, *EEG Rest*, *Pupil Story* | 230 | 7 | Bilateral Amygdala, Bilateral Basal Ganglia, Right Hippocampus |
| *Pupil Rest*, *EEG Rest*, *Movies* | 297 | 6 | Left Amygdala, Bilateral Basal Ganglia, Bilateral Thalamus |
| *Pupil Rest*, *Pupil Story*, *Movies* | 271 | 1 | Right Amygdala |
| *EEG Rest*, *Pupil Story*, *Movies* | 227 | 5 | Bilateral Amygdala, Left Basal Ganglia, Left Hippocampus |
